# Structural and energetic insights into human rhomboid proteases reveal a unique lateral gating mechanism for orphan family members

**DOI:** 10.1101/2025.11.21.689725

**Authors:** Bryony RJ Clifton, Robin A Corey, Adam G Grieve

## Abstract

Rhomboid proteases play fundamental roles in human biology and disease by cleaving substrate transmembrane domains. However, the absence of structural data and the inability to identify substrates for orphan human rhomboids limit mechanistic understanding. Lateral gating is considered the primary route by which transmembrane substrates access the rhomboid active site, although this has been demonstrated directly only for the bacterial rhomboid GlpG. To address this, we characterised the structures, conformational ensembles, and energy landscapes of human rhomboid proteases using AI-based structural models and molecular dynamics simulations, benchmarked by simulations seeded with GlpG crystal structures and models. We find that while some human rhomboids readily transition to open conformations compatible with transmembrane substrate access, orphan rhomboids possess unusually narrow lateral gates that require substantially higher energy to open. Our results reveal unexpected diversity in substrate engagement mechanisms among human rhomboids, and provide a rationale for the orphan status of recently evolved family members.

## Introduction

Intramembrane proteolysis is a fundamental regulatory mechanism that governs the activation and turnover of membrane proteins, with profound implications for cellular signalling and human disease (*1, 2*). The unique structural arrangement of well characterised intramembrane proteases such as bacterial GlpG and human gamma-secretase reveal that their polar active site, which is responsible for the hydrolysis of substrate transmembrane domains within lipid bilayers, is shielded from the hydrophobic membrane environment (*3–6*). This general architecture enables hydrolysis of transmembrane domains but raises a central mechanistic question: how do membrane-embedded substrates gain access to the buried catalytic site?

The rhomboid class of intramembrane proteases is uniquely present in all kingdoms of life, and is characterised by an active site serine, which acts as the nucleophile during proteolysis following abstraction of a proton by the active site histidine (*4, 7*). Across evolution, they share a highly conserved transmembrane fold comprising six transmembrane helices (TMs), termed the rhomboid fold. For substrate to be cleaved, it must enter the rhomboid fold to access the active site. The prevailing model for transmembrane substrate access to rhomboids is lateral gating, which proposes that substrates enter within the plane of the lipid bilayer between TM2 and TM5 of the rhomboid fold (*8*). The bacterial rhomboid GlpG has served as a paradigm for understanding this process, and its study has led to a model in which substrates are first sampled within the rhomboid fold prior to their cleavage. Critically, the physical properties of transmembrane segments determine substrate status. Upon exiting the lipid bilayer and entering the rhomboid fold, it is thought that unstable transmembrane domains (such as those containing helix-destabilising residues) partially unwind, allowing deeper penetration into the rhomboid fold and access to the catalytic dyad. Under the same conditions, transmembrane segments that are helically stable diffuse back into the membrane (*9–11*). This general concept is thought to be true for all intramembrane proteases (*12, 13*). Consistent with lateral gating as a model for substrate access, mutations that promote lateral gate opening in the bacterial rhomboid GlpG enhance proteolysis, whereas mutations that stabilise the closed state inhibit cleavage (*8, 14, 15*). These data collectively demonstrate that an open lateral gate is a prerequisite for substrate entry and cleavage, although it remains unclear whether substrates actively induce gate opening or instead exploit pre-existing conformational dynamics.

Despite these insights from bacterial systems, the extent to which lateral gating governs transmembrane substrate access in human rhomboid proteases remains unresolved. No experimentally determined structure of a human rhomboid protease is currently available (*2, 16*). The degree to which they are functionally similar to GlpG is therefore unknown (*5, 17*). While topological conservation of the core rhomboid fold and the catalytic dyad suggests a shared mechanism, differences in amino acid composition and substrate repertoire raise the possibility of functional divergence within the rhomboid proteases. As dysregulation of rhomboids is linked to a variety of human diseases, including Alzheimer’s and Parkinson’s Disease, it is important that we gain mechanistic insights into rhomboid intramembrane proteolysis in humans (*18–20*).

In mammals, rhomboids can be classified into secretase and mitochondrial proteases. The secretases are rhomboid-like (RHBDL) 1-4, and the mitochondrial rhomboid is presenilin-associated rhomboid-like (PARL) (*16, 21*). As described earlier with GlpG, the core rhomboid fold has six TMs. RHBDL4 is predicted to have only these six TMs, while RHBDL1-3 have an extra C-terminal domain (designated 6+1), and PARL has an extra N-terminal domain (designated 1+6). PARL is best known for its role in mitochondrial quality control through cleavage of PINK1, a kinase frequently mutated in Parkinsons Disease (*20, 22, 23*). The best characterised mammalian secretase rhomboid, RHBDL2, has several known substrates including single-pass thrombomodulin, ephrin-Bs, adhesion proteins, EGFR, and the multi-pass membrane protein Orai1 (*10, 24–26*). As a result, RHBDL2 has been implicated in wound healing, cell-cell adhesion, and T cell activation. RHBDL4, like PARL, has also been shown to perform quality control functions, amongst other roles. For instance, its known substrates include pre-T cell receptor (pTα), TMED7 and APP, as well as soluble proteins within the lumen of the endoplasmic reticulum (ER). Accordingly, RHBDL4 is implicated in ER-associated degradation, inflammatory signalling, and Alzheimer’s Disease (*27–31*). The other two mammalian rhomboids, RHBDL1 and RHBDL3, are referred to as the orphans of the family, as no substrates have been identified for them, and so have no ascribed cellular roles. Their conserved active site residues are accessible to soluble activity-based probes, which indicates that they are indeed proteases (*32*). However, as the orphans arose more recently in metazoans, they may have novel roles and biomechanics. They are enriched in the nervous system, and RHBDL3 is linked to ageing and neurological disease; therefore identifying the biological role of orphan rhomboids is crucial for our understanding of these processes (*33, 34*).

We questioned whether advances in structural and conformational prediction tools combined with *in silico* approaches for exploring biomolecular mechanics could shed light on the function of human rhomboid proteases. AI-based structure prediction tools such as AlphaFold, Boltz, and Chai have revolutionised our ability to obtain insights into 3D biomolecular structures, especially for membrane proteins that are challenging to study using traditional experimental methods (*35–37*). However, structural predictions for membrane proteins and their membrane-based interactions (for instance in AlphaFold Multimer mode) can be unreliable as they are generated without explicitly accounting for the native membrane environment (*38*). A recent analysis revealed ambiguity and a lack of confidence in many membrane protein structure predictions (*39*). These structural predictions therefore require validation through molecular dynamics (MD) simulations within artificial membrane systems, which allow for capture of realistic conformations and interactions.

Here, we combine AI-based structural prediction with physics-based MD simulations to investigate the mechanisms of human rhomboid proteolysis. By modelling rhomboid proteases in the presence and absence of substrates, and simulating their behaviour within lipid bilayers, we assess both structural plausibility and dynamic function. Our results reveal that the lateral gating mechanism identified for GlpG is broadly conserved in human rhomboid proteases. We propose a model in which stochastic lateral gate opening is coupled to substrate conformational sampling, with productive cleavage arising from the temporal alignment of gate opening and substrate helix destabilisation. Notably, however, the orphan rhomboid proteases RHBDL1 and RHBDL3 represent an anomaly in this landscape: each exhibit a narrower lateral gate and altered energetic requirements for substrate access. These features point to a potential evolutionary adaptation involving conformational regulation or specialised substrate recognition, which are likely required to overcome the energetic barrier at the orphan lateral gate. Elucidating these mechanisms will be essential for understanding the roles of the orphan rhomboids in ageing and neurological disease.

## Results

### GlpG spontaneously transitions between open and closed states

The study of GlpG has established a general structural and functional template for rhomboid proteases. Structural analyses of GlpG have shown that transmembrane helices TM2 and TM5 function as a lateral gate that regulates substrate access to the serine-histidine catalytic dyad (*2, 8, 14, 17*). To assess the reliability of AI-derived structural predictions of rhomboid proteases, we benchmarked GlpG structural predictions against crystal structures of *E.coli* GlpG in different conformational states. The open (Fig. 1A; PDB: 2NRF) and closed (Fig. 1B; PDB: 2IC8) crystal structures of GlpG highlight pronounced differences in lateral gate positioning. Measurement of the average distance between two residue pairs positioned in the plane of the active site revealed a lateral gate width of approximately 1.09 nm in the closed state, and 1.55 nm in the open state. Because any experimentally determined structure represents a qualitative snapshot within a potentially broader conformational landscape, we performed 5 x 2 µs atomistic MD simulations to investigate the intrinsic conformational dynamics of GlpG. In particular, we were curious as to whether simulations seeded with the closed structural model could spontaneously transition to a state consistent with substrate gating, or whether such transitions required substrate. Simulations of both crystal structures demonstrated that GlpG can readily transition between open and closed states in the absence of substrate or inhibitors (Fig. 1C, Fig. S1 and S2), indicating it can naturally sample a dynamic ensemble of conformations. Notably, over the course of the simulations, the closed state of GlpG (2IC8) sampled substantially wider lateral gate conformations, approaching widths comparable to that observed in the original open state structure (2NRF; ∼ 1.55 nm), which itself sampled even wider lateral gate states. This suggested that even in the absence of substrate the rhomboid lateral gate can spontaneously transition from a closed to an open conformation, capable of accommodating substrate.

**Fig. 1-.**
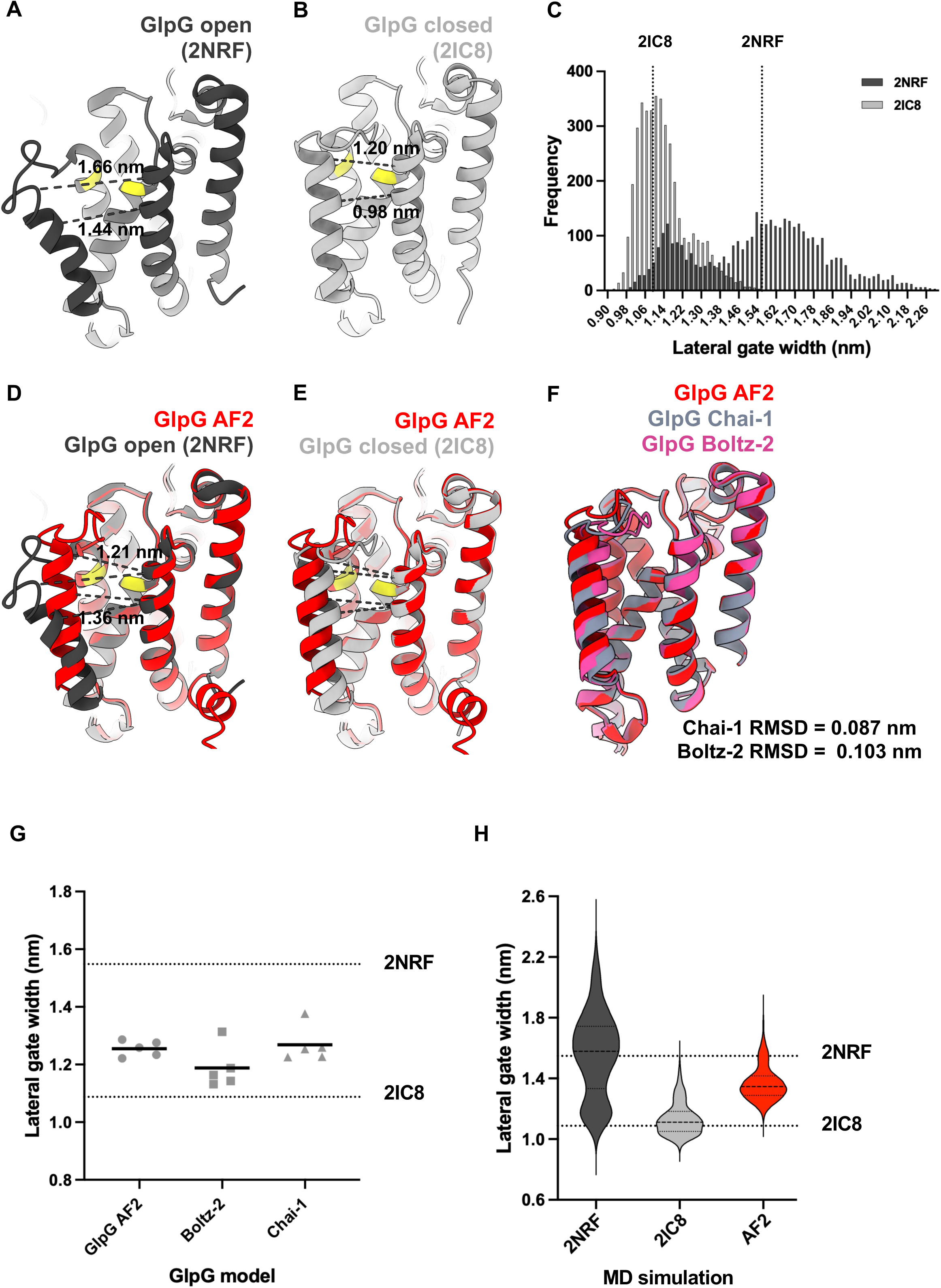
GlpG spontaneously transitions between closed and open states. **(A-B)** Cartoon representations of E. coli GlpG structures 2NRF (A) and 2IC8 (B). Lateral gate helices (TM2 and TM5) are labelled. The active site is shown in yellow and dashed grey lines show the residue pair above and below the plane of the active site, which was used to used to calculate the average lateral gate width (alpha-carbon atoms of M149-G240 and N154-W236). (C) Histograms showing the overlaid distributions of lateral gate widths across five 2 μs simulations of equilibrium simulations in a 7:3 POPC:cholesterol lipid bilayer, seeded with either 2NRF or 2IC8 structures as indicated. Dashed lines indicate the lateral gate distance of 2NRF and 2IC8 crystal structures. **(D-E)** Structural alignments of PDB ecGlpG crystal structures with the GlpG AF2 rank 1 model. (**F**) Structural alignment of the core fold from ecGlpG rank 1 models from AF2, Chai-1, and Boltz-2. RMSD values for the core fold, relative to the AF2 model, are indicated. (**G**) Lateral gate quantification of the top 5 ranked ecGlpG models generated from diverse structure prediction tools. Black bar = mean. Dashed line = crystal structure lateral gate width. (**H**) Violin plots showing the lateral gate distributions of 2NRF, 2IC8 and AF2 MD simulations of ecGlpG. Dashed lines indicate the post-equilibrium lateral gate width of models used to seed simulations.

We next evaluated whether current AI-based structural prediction tools, such as AlphaFold, Chai-1 and Boltz-2, generate models consistent with these experimentally observed parameters. AF2 produced structures with lateral gate distances that fall between the open and closed crystal structures (Fig. 1D-E). By comparison, Chai-1 and Boltz-2 produced very similar models and lateral gate widths that sat between the open and closed crystal structures (Fig. 1F-G, Supplementary Table 1). Simulations of the highest ranking AF2-predicted model revealed similar conformational flexibility to the experimentally determined structures, with the GlpG lateral gate sampling a range of approximately 1.1 nm to 1.8 nm (Fig. 1H). Together, these data demonstrate that GlpG inherently samples a broad spectrum of conformational states and is energetically accessible to transmembrane domains. These data also show that AI-based structural prediction approaches yield models that are consistent with experimentally determined conformations.

### Structural predictions provide new insights into the mammalian rhomboid proteases

As there are no experimentally-solved structures for any mammalian rhomboid protease, it is unclear whether they are mechanistically similar to GlpG. We therefore employed a suite of structural prediction tools and assessed the quality of the generated models (Fig. 2, Fig. S3 and S4; see Supplementary Tables 1 and 2 for quality metrics; for structural models see https://osf.io/n6348/) (*40, 41*). Overall, confidence predictions for the AF2 models of human rhomboids were highest for RHBDL2 and PARL (0.84 and 0.85 pTM respectively for rank 1), indicating good predictions (Fig. S4A). RHBDL1 and RHBDL3 models were lower confidence, although low confidence regions were generally confined to interhelical loops, or cytoplasmic/IMS domains, which are likely highly flexible, as reflected in pLDDT scores (Fig. S4, B and D). RHBDL4 had the lowest pTM score (0.64), largely due to a highly unstructured C-terminus (Fig. S4E). These cytoplasmic domains (the IMS domain in PARL) reflect a striking and expected point of heterogeneity between the different rhomboids. For all systems, the pLDDT scores of the core rhomboid fold were high (Fig. S4, B-G), indicating high confidence in the models at this region. The confidence of the core rhomboid fold was also evident from evolutionary coupling analysis, which revealed strong co-evolution among transmembrane residues, supporting the accuracy of the structural prediction (Fig. S5) (*42*). Furthermore, sites of evolutionary coupling closely match those of 3D contacts within the core rhomboid fold (Fig. S6).

**Fig. 2-.**
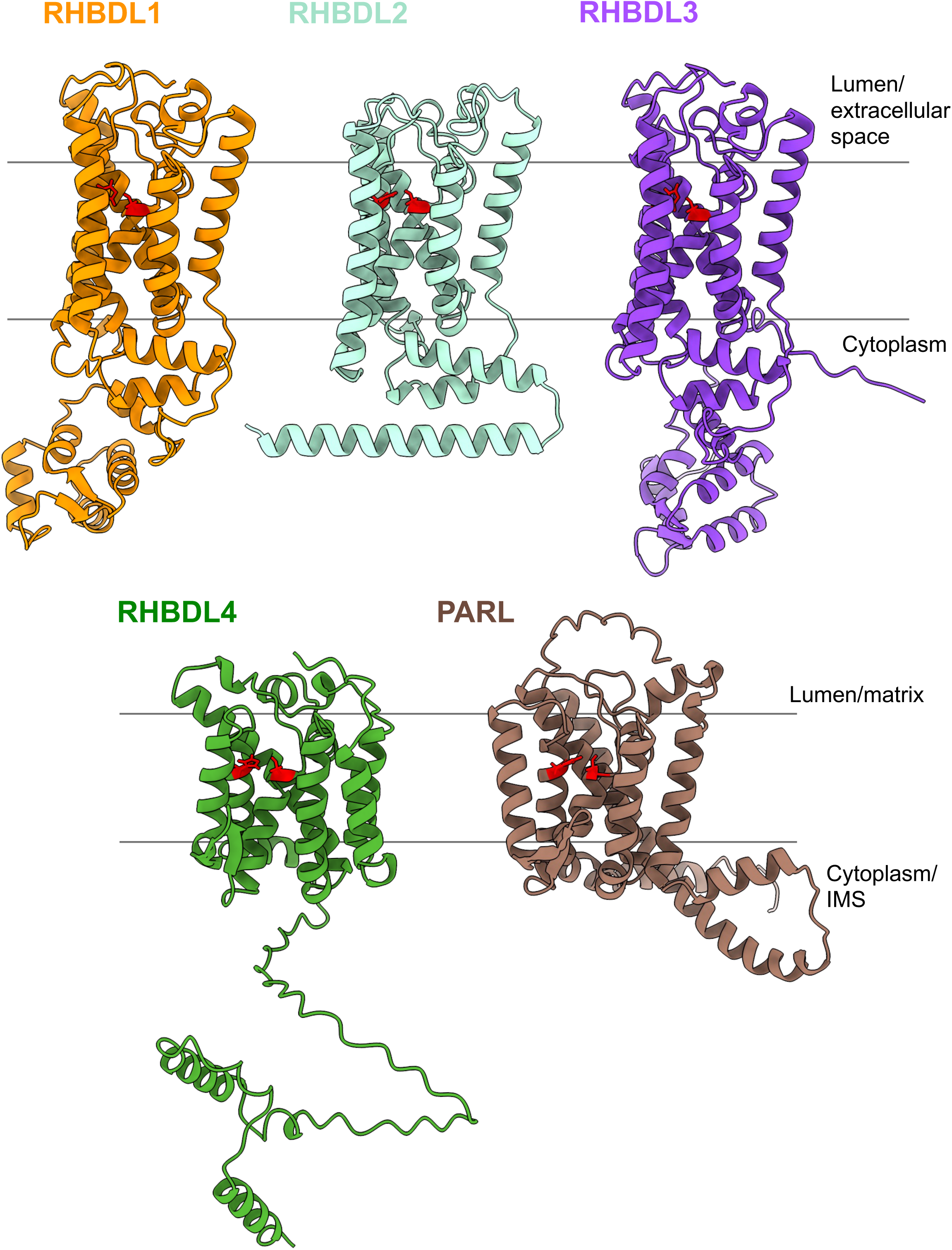
AlphaFold2 predictions of mammalian rhomboids. Cartoon representations of rank 1 models for each of the human rhomboid proteases. Active site residues are in red. Core rhomboid folds are similar but soluble domains show high heterogeneity. For RHBDL1-3, the cytoplasmic domain is N-terminal, whereas this is C-terminal for RHBDL4 and PARL due to their 1+6 topology in the models. Additionally, RHBDL1-3 possess two short amphipathic helices between the core fold and the cytoplasmic domain, whereas RHBDL4 and PARL appear to lack these. Whereas the N-terminal domain of RHBDL2 is a single, continuous helix, for RHBDL1 and -3, this domain is extended and folded into a folded domain comprising two EF-hand motifs (see Fig. S16). In contrast, the cytoplasmic domain is predicted to be largely unstructured for RHBDL4. For PARL, this matrix domain is the end of one continuous helix extending down from TM6 and terminating in a small unstructured region.

Hydrophobic mismatch with the membrane has been found to be central to rhomboid function (*43, 44*); accordingly, all TMs within each rhomboid were observed of different lengths (Fig. S4H). Most models of the rhomboids matched general structural expectations of having seven transmembrane helices, with RHBDL1-3 exhibiting a 6+1 topology, and PARL presenting with a 1+6 topology. However, the AF2 model for RHBDL4 has a 1+6 topology similar to PARL, which contrasts with previous studies indicating it has only six transmembrane domains (*27*). Interestingly, we noticed that the rhomboid fold TMs in RHBDL1-3 appear to have very similar lengths, whereas RHBDL4 and PARL harbour many short TMs which may mirror the membrane thickness of the subcellular compartments to which they target (Fig. S4H). These results indicate an unanticipated similarity between RHBDL4 and PARL (Fig. S7A-D), which is discussed later. Overall, the differences in structures of the human rhomboids mirror their evolutionary relationships, with RHBDL1-3 being most closely related, and PARL and RHBDL4 being divergent from other members of the family (*45–47*).

### AF2 models reveal lateral gate differences between characterised and orphan rhomboids

We next examined the lateral gates across the different structural models. AF2 predictions for the human rhomboids revealed a striking degree of heterogeneity in lateral gate width. Top-ranked AF2 models for each protein were structurally aligned to RHBDL2, the best characterised secretase rhomboid in humans. Notably, whilst the RHBDL2 lateral gate is narrower than that of GlpG (Fig. 3A-B; 1.06 nm and 1.26 nm respectively), it is even more constricted in the two orphan rhomboids relative to their closest homologue (Fig. 3C-D; 0.99 nm and 0.96 nm respectively). Indeed, the lateral gates in RHBDL1 and RHBDL3 were narrower than in all other rhomboid models, even the closed crystal structure of GlpG (ca. 1.09 nm). In contrast, the lateral gates of RHBDL4 (1.19 nm) and, to a greater extent, PARL (1.76 nm) are substantially wider than that of RHBDL2 (1.06 nm), indicating an unexpectedly broad spectrum of lateral gate width across the human proteases (Fig. 3E-F). As has been reported for GlpG, lateral gate helices and connecting loops show low confidence in nearly all human rhomboid predictions, suggesting they are dynamic regions across the family (Fig. S4) (*14*).

**Fig. 3-.**
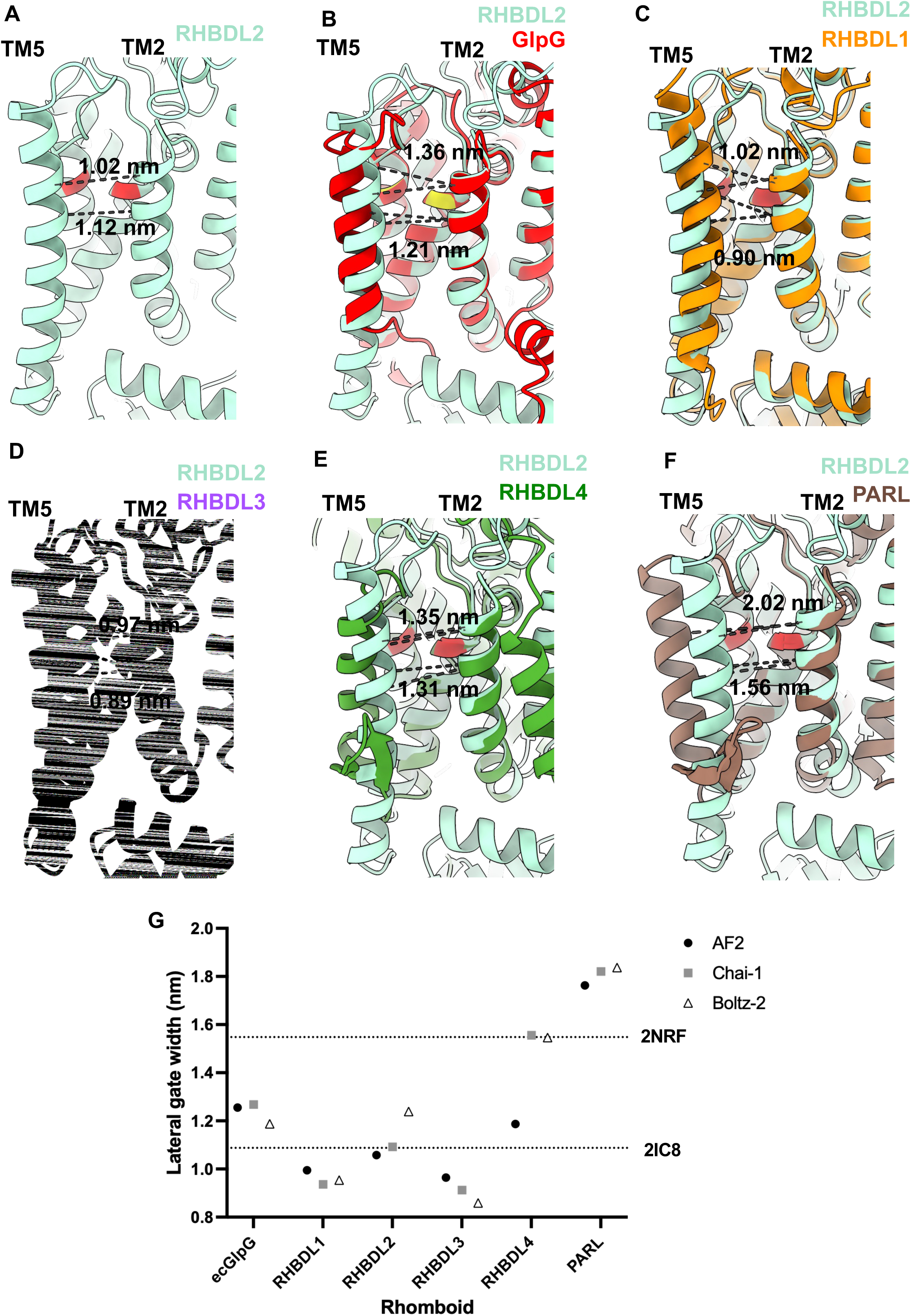
Lateral gate measurements for AlphaFold2 predictions of human rhomboids and *E. coli* GlpG. Equivalent residue pairs in TM2 and TM5 were used for lateral gate quantification of ecGlpG and the human rhomboids. (measurements indicated with dashed line). Images show cartoon representations of the rhomboids, looking through the lateral gate into the active site (yellow for GlpG, red for the human rhomboids). Lateral gate helices (TM2 and TM5) are labelled. **(A)** RHBDL2, **(B-F)** Structural alignments of GlpG (B) and human rhomboids to RHBDL2 (C-F). **(G)** Plot of lateral gate distances for the top-ranking model of each rhomboid, generated with AF2, Chai-1, or Boltz-2. For comparison, the lateral gate width for the solved structures of GlpG in its closed (2IC8) and open (2NRF) conformation as depicted by dotted lines.

Models produced by AF2, Chai-1 and Boltz-2 consistently indicate that this variability is a robust and reproducible feature of the predictions, rather than a model-specific artefact (Fig. 3G). The only exception are the RHBDL4 models, which displayed substantial heterogeneity both between tools and within individual prediction sets, in line with its comparatively lower confidence metrics (Supplementary Table 1). This variability may reflect the capacity of AI-driven structural prediction methods to capture multiple biologically relevant conformational states, as reported previously, and indicates that the RHBDL4 rhomboid fold is intrinsically more flexible than that of other rhomboids (*48*). Consistent with this idea, a shared beta-sheet structure between TM4 and TM5 is evident in the RHBDL4 and PARL models, which may relate to the wider lateral gate conformations (Fig. S7D).

In summary, structural models of human rhomboids generated using diverse AI-based approaches reveal distinct features of the orphan rhomboids relative to their better characterised counterparts. Saliently however, none of the apo models for RHBDL1-3 appear to adopt fully gated or ‘open’ conformations, when benchmarked against the lateral gate width of the ‘open’ state of GlpG (1.55 nm) (Fig 3G).

### AF2 supports a substrate gating mechanism for mammalian rhomboids

Our recent work revealed that the full-length form of the substrate PINK1 can be modelled within the rhomboid fold of PARL with AF2 (Fig. 4A) (*49*). In this model, the lateral gate adopts a width of ∼1.8 nm, consistent with that observed across all apo models of PARL (Fig. 3G). Notably, all PARL models exhibit lateral gates that are significantly wider than any of our apo models of the secretase rhomboids. We therefore reasoned that the lateral gate widths measured in the apo models of secretase rhomboids (Fig. 3G) are prohibitively narrow for transmembrane substrate entry, and so must be capable of widening significantly further. To test this hypothesis, we assessed whether AF2-multimer could model entry of known substrates into the lateral gate of human secretase rhomboids. Specifically, we generated predictions for RHBDL2 and RHBDL4 in complex with the isolated transmembrane segments of known substrates, either Orai1 TM4 or pTα respectively (Fig. 4B-C). The resulting structural models of substrate gated RHBDL2 and RHBDL4, in which the substrate is positioned adjacent to the rhomboid active site, provide strong support for a mechanism in which the human rhomboid proteases access transmembrane substrates via lateral gating between TM2 and TM5.

**Fig. 4-.**
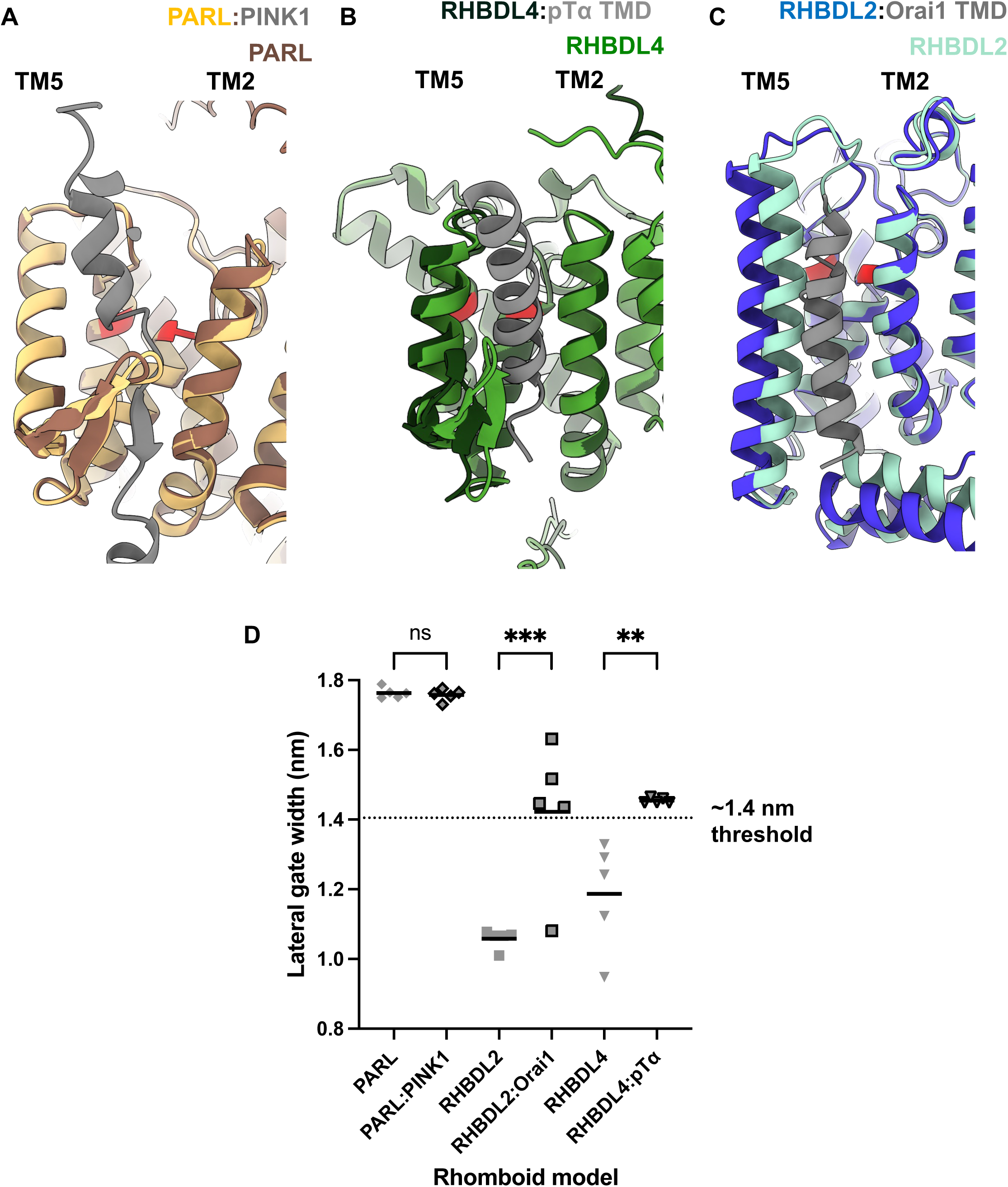
AlphaFold2 predicts substrate gating for human rhomboids. **(A-C)** Overlays of structurally aligned rank 1 AF2 predictions of apo human rhomboids and substrate-bound human rhomboids as labelled. **(D)** Quantification of lateral gate widths for rhomboids either with or without substrate. Dashed lines indicate the threshold for LG width associated with gating of substrate (1.4 nm), which is the width in the rank 1 model of RHBDL2:Orai (post equilibration in a lipid bilayer). Bar = mean value. One-way ANOVA with Šídák’s multiple comparisons test. One asterisk represents P < 0.033; two asterisks represent P < 0.002; three asterisks represent P < 0.001.

In line with our hypothesis, the lateral gates of both RHBDL2 and RHBDL4 significantly widened in the presence of substrate (1.06 nm to 1.42 nm for RHBDL2 apo vs substrate gated; 1.19 nm to 1.45 nm for RHBDL4 apo vs substrate gated) (Fig. 4D). This was in stark contrast to the PINK1:PARL model, where the lateral gate did not widen compared to the apo models, suggesting that AF2 captures PARL in a fully open state, both in the presence and absence of substrate. RHBDL4 shows a similar, but more heterogeneous profile to PARL – with a modest widening of the lateral gate in the presence of substrate, relative to RHBDL2. As both RHBDL4 and PARL perform protein quality control functions, this may reflect their ability to constitutively sample the membrane proteome of their respective organelles. Overall, these data highlight that for human rhomboids to gate substrates, their lateral gates must open to ∼1.4 nm – just wide enough to fit an α-helix (diameter ca. 1.2 nm).

### MD simulations reveal that the lateral gate of characterised rhomboids can transition to open conformations in the absence of substrate

We next questioned whether, as observed for GlpG (Fig. 1), the apo forms of human rhomboids are intrinsically conformationally flexible and thus primed for substrate entry. To address this, we employed atomistic MD simulations to first test the stability and validity of the models within a membrane bilayer environment, and subsequently characterised their conformational dynamics. All rhomboids were simulated in identical lipid bilayer environments to enable direct comparison. Route mean square deviation (RMSD) analysis was performed to evaluate structural stability. Notably, the cytoplasmic domains of all secretase rhomboids displayed unusually elevated flexibility throughout simulations, therefore additional RMSD calculations were performed in their absence (Fig. S8-9). As the simulations converged to stable RMSD values, typically below 0.3 nm, the AF2 predictions appear to successfully capture energetically favourable conformations, supporting their validity.

We next quantified lateral gate width over the course of the simulations to assess the relative dynamics of TM2 and TM5 in each of the rhomboids, focussing initially on the well characterised RHBDL2, RHBDL4 and PARL (Fig. 5A-B, S9). This analysis revealed that RHBDL2 samples a broad distribution of lateral gate conformations (Interquartile range/IQR 1.07-1.37 nm across the five runs, Fig. 5A,C-D), comparable to that observed in GlpG simulations (1.29-1.42 nm, Fig 1H). Notably, RHBDL2 frequently exceeds the ∼ 1.4 nm threshold associated with substrate gating (Fig. 4 C-D; 21.3% simulation time above threshold, similar to GlpG for which 29.5% of simulation time is above 1.4 nm). This indicates that apo GlpG and RHBDL2 can readily access conformations permissive for transmembrane substrate entry.

**Fig. 5-.**
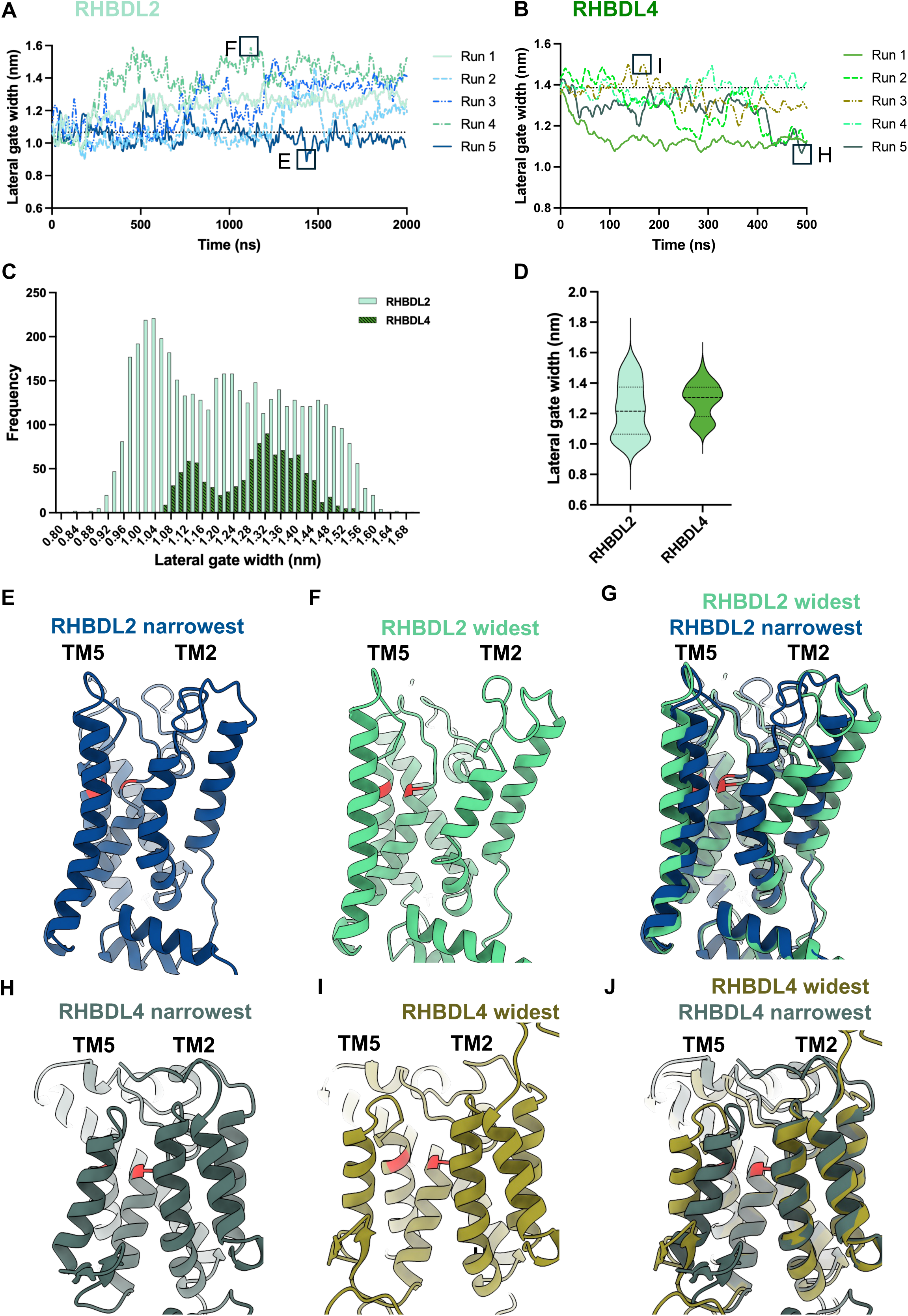
Molecular dynamics simulations of characterised secretase rhomboids RHBDL2 and RHBDL4. Lateral gate distance profiles for **(A)** RHBDL2 and **(B)** RHBDL4 during molecular dynamics simulations. Rank 1 AF2 models were used as inputs following generation of a 7:3 POPC:cholesterol lipid bilayer and energy minimisation. Five 2 μs and five 500 ns simulations were performed for RHBDL2 and RHBDL4 respectively. Each simulation is shown as a separate line. Dashed line shows the input lateral gate width (post equilibration in a lipid bilayer). **(C)** Histograms show the lateral gate width distributions across simulations shown in (A) and (B). **(D)** Violin plots comparing RHBDL2 and RHBDL4 lateral gate width profiles**. (E, F)** Snapshots of RHBDL2 narrowest and widest states from (A). **(G)** Overlay of structural alignments of (E) and (F). **(H, I)** Snapshots of RHBDL4 narrowest and widest states from (B). **(J)** Overlay of structural alignments of (H) and (I).

RHBDL4 similarly explores a wide conformational landscape. However, in contrast to RHBDL2, its lateral gate often narrows from its initial open state (∼ 1.4 nm) across multiple trajectories (Fig. 5B-D) (IQR 1.18-1.37 nm). Notably, RHBDL4 occupies an lateral gate conformation frequently (16.5% of simulation time above 1.4 nm). Together, these findings indicate that both RHBDL2 and RHBDL4 have dynamic lateral gates capable of transitioning between open and closed conformations with relatively low energetic cost. Consistent lateral gate dynamics were also observed for RHBDL4 using a more ER-like lipid composition (Fig. S10), suggesting that these behaviours arise from intrinsic features of the RHBDL4 fold rather than the membrane environment.

In contrast ro RHBDL2 and RHBDL4, the lateral gate of PARL remains persistently open throughout simulations (IQR 1.65-1.76 nm), sampling almost an entirely distinct ensemble from RHBDL2 or RHBDL4 (Fig. S10), pointing to a distinct mechanism for substrate access.

Overall, these data suggest that, like GlpG, both RHBDL2 and RHBDL4 dynamically interconvert between open and closed states at equilibrium with minimal energetic cost, albeit with different conformational preferences: RHBDL2 favouring more closed states, RHBDL4 more open states (Fig. 5D), and PARL being constitutively open. Representative snapshots of the narrowest and widest states are shown in Figure 5E-J. Collectively, our data reveal that the lateral gates of RHBDL2 and RHBDL4, both of which have known substrates, can transiently adopt stable conformations compatible with substrate entry.

### MD simulations reveal that orphan rhomboids favour closed lateral gate conformations

Given that MD simulations of the apo forms of RHBDL2 and RHBDL4 revealed frequent sampling of open lateral gate conformations (Fig. 5), we next asked whether the same was true for the orphan secretase rhomboids. However, in contrast to their better-characterised secretase counterparts, MD simulations initiated from the AF2 models for RHBDL1 and RHBDL3 show that their lateral gates remain comparatively narrow (IQR 0.83-1.06 nm and 0.90-1.07 nm respectively), and – in the timescale of our simulations – rarely cross the ∼1.4 nm threshold associated with substrate entry (Fig 6; simulation time above this threshold is 1.9% for RHBDL1 and <0.1% for RHBDL3). This behaviour is consistent with the AF2 models representing energy minima (Fig. 3C-D; Fig. 6A-B). Although the lateral gates of both RHBDL1 (0.90 nm starting width), and RHBDL3 (0.88 nm starting width) briefly sample states consistent with gating during the simulations they generate similar conformational profiles that are clearly distinct from those of RHBDL2 and RHBDL4 (Fig. 6C). Across all human rhomboids, RHBDL1 and RHBDL3 maintain significantly narrower lateral gates (Fig. 6D). As rhomboid lateral gate opening is required for substrate access, these data may explain why RHBDL1 or RHBDL3 substrates have not been readily identified. Despite their typically narrower lateral gates, these simulations reveal that the orphans can sample open states in the absence of substrate, but they do so infrequently. Figure 6E-J illustrates the range of lateral gate conformations the orphan rhomboids can occupy with snapshots of the narrowest and widest states sampled in MD simulations (Fig. 6E-J).

**Fig. 6-.**
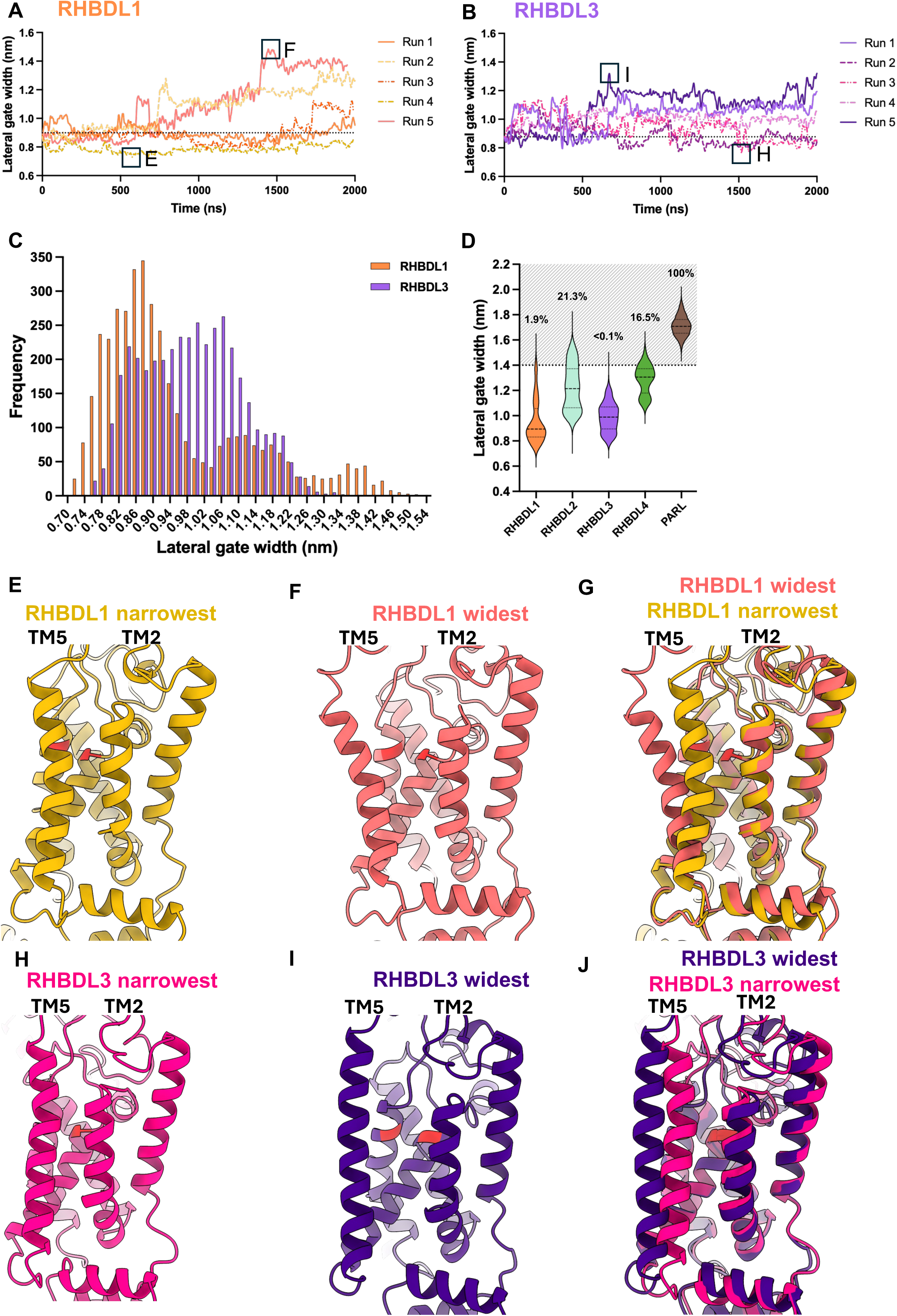
Molecular dynamics simulations of orphan secretase rhomboids show stably narrow lateral gates. Lateral gate width profiles for **(A)** RHBDL1 and **(B)** RHBDL3 during molecular dynamics simulations. Rank 1 AF2 models were used as inputs following generation of a 7:3 POPC:cholesterol lipid bilayer and energy minimisation. For each rhomboid, five 2 μs simulations were performed. Each simulation is shown as a separate line. Dashed lines show the input lateral gate width (post equilibration in a lipid bilayer). **(C)** Histograms show the lateral gate width distributions across simulations shown in (A) and (B). **(D)** Violin plots comparing lateral gate width profiles of all human rhomboid simulations (Including data from Fig. 5 and Fig. S9). The dashed line indicates the ∼1.4 nm gating threshold and the cumulative frequency of simulation time above this threshold is displayed above each density curve as a percentage. **(E, F)** Snapshots of narrowest and widest states of RHBDL1 from (A). **(G)** Overlay of structural alignments of (E) and (F). **(H, I)** Snapshots of narrowest and widest states of RHBDL3 from (B). **(J)** Overlay of structural alignments of (H) and (I).

To further characterise the differences between orphan rhomboids and those with known substrates, we performed a principal component analysis (PCA) on the simulation trajectories, to determine the extent to which lateral gate motions contribute to overall dynamics (Figure S11). RHBDL1-3 were selected due to their highly conserved core fold, including having near identical TM lengths (Fig. S4H). Performing PCA across all apo trajectories revealed that the first eigenvector accounted for ∼41% of the total variance in the system (Fig. S11A). Visualisation of the extreme states for this eigenvector revealed that eigenvector 1 mostly represents motions within the lateral gate helices (Fig. S11B). Strikingly, projection of the individual trajectories onto this eigenvector revealed that RHBDL1 and RHBDL3 cluster together, with a slight overlap, whereas RHBDL2 is entirely distinct (Fig. S11C).

Together, these analyses confirm that lateral gating is a dominant feature of rhomboid dynamics and demonstrates, in an unbiased approach, that the orphan rhomboid proteases display distinct conformational ensembles that markedly differ from those of the characterised rhomboid RHBDL2. Importantly, these insights indicate that orphan rhomboids retain the capacity to access catalytically competent conformations, yet such states are only rarely sampled by AF2 predictions or equilibrium MD. A plausible explanation is that lateral gate opening in orphan rhomboids is associated with a higher energetic barrier, limiting its spontaneous occurrence.

### Steered MD and potential of mean force calculations reinforce that energy barriers are associated with lateral gate opening for orphan rhomboids

To address whether a higher energy barrier is associated with lateral gate opening for the orphan rhomboids, steered MD was performed to gradually drive the human secretase rhomboid proteases into an extreme open state (Fig. 7, A-C depicts this with RHBDL3 as an example; Fig S12A shows the distance profile from this simulation). This allowed for a broad spectrum of lateral gate widths to be sampled, beyond those achieved by equilibrium MD. The conformational landscape associated with lateral gate dynamics could then be assessed using umbrella sampling to calculate the potential of mean force (PMF) of LG widening.

**Fig. 7-.**
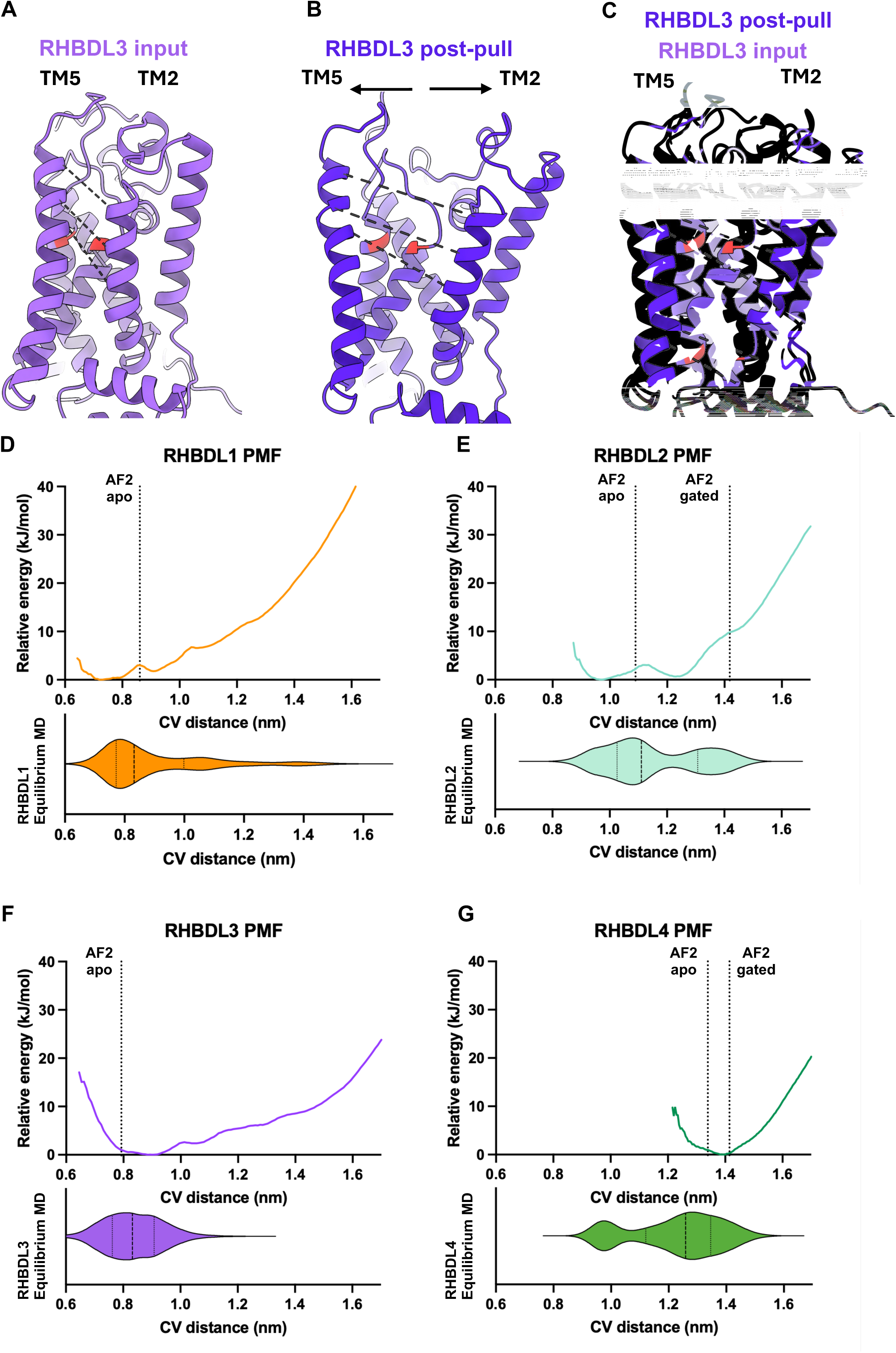
Steered molecular dynamics and energy landscapes associated with gating of secretase rhomboids. Constant velocity steered molecular dynamics were applied to alpha-carbon atoms within 3 residue pairs on TM2 and TM5 (‘collective variable’ or CV), as indicated by dashed grey lines. Following steered molecular dynamics of RHBDL1-4, frames were selected using even spacing of ∼0.05 nm and used to seed umbrella sampling simulations. Potential of mean force (PMF) calculations were performed, allowing an energy landscape associated with lateral gate dynamics to be constructed. **(A)** Cartoon representation of the input model. **(B)** Cartoon representation of the final frame from the pull simulation of RHBDL3 (‘post-pull’). **(C)** Overlay of (A) and (B). **(D-G)** Relative energy landscape for RHBDL1, -2, -3 and -4 respectively (top) and corresponding violin plots showing the distribution of lateral gate widths in equilibrium MD simulations of apo models, using equivalent CV quantification as was used for steered MD and PMFs. Dashed lines indicate lateral gate width for either apo or gated AF2 models (post equilibration in a lipid bilayer).

To steer open the lateral gate, three paired residues along TM2 and TM5 were selected upon which a pulling force was applied (termed the ‘collective variable’ or CV). The position of the CV was selected to ensure uniform pulling across the helices in a manner congruent with the gated or open rhomboid models. It should be noted that, as a result, CV distance measurements vary slightly from lateral gate width measurements in previous analyses, which used two residue pairs in the plane of the active site. Where relevant, these two measurements will both be compared for clarity.

Equilibrium MD simulations seeded from the output of steered MD were performed to evaluate whether the human secretase rhomboids can stably exist in these extreme open states (Fig. S12). As expected, all rhomboids narrowed from this extreme open state. However, both RHBDL1 and RHBDL3 appear to stabilise at an intermediate, gated open state similar to the rare state seen in simulations of AF2 models (Fig. S12B-C). This indicates that orphan rhomboids can stably occupy an open state, which likely represents a local energy minimum.

Following steered MD to force apart lateral gate helices, the free energy landscape associated with the conformational change (the PMF) was explored by selecting frames at regular intervals throughout these pull simulations. These frames were used to seed umbrella sampling simulations, which apply a biasing potential to the CV, acting as an energetic penalty to deviations from the starting conformation within each window. A relative free energy landscape was then constructed using the Weighted Histogram Analysis Method (WHAM) to reweight the data and correct for any biasing introduced during the umbrella sampling (*50*). Energy landscapes associated with lateral gate dynamics were built for the secretase rhomboids RHBDL1-4. In these landscapes, wells represent local minima, or low energy protein conformations. Umbrella histograms show that all states were sufficiently sampled (Fig. S13). For comparison, we have super-imposed the lateral gate widths from AF2 predictions of apo, and where available, gated rhomboid models onto energy landscapes (Fig 7D-G, dashed lines). Violin plots are also shown for corresponding AF2-MD equilibrium simulations, using the same three paired residues along TM2 and TM5 used for CV quantification as a reference (data replotted from Figs. 5 and 6).

The energy landscapes show that the RHBDL1 lateral gate is the narrowest of the human rhomboids with a primary energy well seen at ∼0.7 nm (Figure 7D). Applying this CV distance quantification to the original AF2 MD simulations shows that this agrees well with the equilibrium MD data (ca. 0.8 nm; Fig. 6A,C-D). A pair of smaller energy wells are also seen at wider states, e.g. at ∼0.9 nm and ∼1.1 nm, supporting a conclusion that we have sampled a rare conformational state above 1.4 nm. For RHBDL2, two primary wells and one smaller energy well are observed, at CV distances of ∼0.95 nm, ∼1.25 and ∼ 1.45 nm (Fig. 7E). These wells represent the different states for RHBDL2 sampled in equilibrium MD simulations (ca. 0.9-1.1 nm, 1.3 nm and 1.5 nm; Fig 5C). The two primary wells are associated with very similar relative free energies and there is a small energy barrier of ca. 3.1 kJ/mol (approximately 1.5 kT) between them. The primary RHBDL3 well is very broad, with a relatively flat region between 0.8-0.95 nm (Fig. 7F). This corresponds well with the equilibrium MD data which shows that RHBDL3 can freely sample a range of states between 0.7-1.0 nm but rarely reaches a conformation compatible with transmembrane substrate entry. One well is seen for RHBDL4 between ∼1.3-1.4 nm (Fig. 7G), matching AF2 predictions and equilibrium MD data (ca. 1.2-1.4 nm; Fig. 5C-D). However, as the rank 1 RHBDL4 AF2 model was in a more open conformation than those of RHBDL1-3, its narrower states were not sampled. Presumably, an additional small well would be seen at ca. 1.0 nm were the landscape to be extended downwards.

In summary, our analysis of AI-based models of human rhomboids reveals that lateral gate distances wider than ∼1.4 nm (a distance consistent with substrate gating, Fig. 4) can be sampled by all human rhomboid proteases (Fig. 5-7). RHBDL4 and PARL appear to be able to enter these states with relatively little energetic cost, however the same is not true for RHBDL1-3. Significant energy wells around and above 1.4 nm are rarely seen in RHBDL1-3 landscapes. For RHBDL2, a small energy well can be seen, but at a much greater relative kJ/mol than for RHBDL4, suggestive that these open states are energetically unfavourable without something to stabilise them (discussed later). Accordingly, MD simulations using gated RHBDL2 from AlphaFold-Multimer predictions demonstrate that once this energy barrier is crossed, RHBDL2 stably remains in this open state (Fig. S14). Further supporting this point, the gated RHBDL2 simulations occupy an open conformation for a much higher proportion of the simulation duration than the apo form (62.0% of simulation time is above 1.4 nm gating threshold for the gated, compared to 21.3% for the apo). These findings mirror our equilibrium MD results with apo rhomboids, where lateral gate widths >1.4 nm are frequently observed with RHBDL4, rarely observed with RHBDL2 and hardly observed with the orphan rhomboids (Fig 7D-G, lower violin plots).

Overall, these data reveal that orphan rhomboids have the capability to gate substrates, but compared to all other human rhomboids, higher energy barriers restrict them from occupying open states.

## Discussion

The present work provides a mechanistic framework for understanding the molecular mechanisms of rhomboid intramembrane proteolysis in humans by addressing the central question of how substrates gain access to a buried catalytic site within the membrane. By integrating AI-based structural prediction with MD simulations, we show that lateral gating is a conserved feature of rhomboid proteases, but one that is differentially tuned across the human rhomboids. These differences are best understood not simply in structural terms, but through the lens of their underlying conformational energy landscapes.

Using GlpG as a benchmark (Fig 1), we demonstrate that human rhomboids sample a continuum of lateral gate conformations (Fig. 5-6), including those compatible with substrate entry (Fig. 4). Importantly, AI-derived models recapitulate experimentally observed states and, when combined with MD simulations, occupy energetically favourable conformations. Across all rhomboids assessed, TM5 was consistently responsible for the majority of lateral gate dynamics, with RMSF analyses showing that TM4/5 and TM5/6 loops were hotspots of flexibility (Fig. S15). This is in agreement with previous observations indicating that an outward movement of TM5 drives substrate gating (*5, 8, 14, 51, 52*). However, rather than a uniform mechanism, our data reveal that individual rhomboids occupy distinct conformational space, indicting that lateral gating is functionally tuned for diverse protein function.

A clear stratification emerges across the human rhomboid proteases. RHBDL2 and RHBDL4 readily sample closed and open lateral gate conformations at equilibrium, with relatively low energetic barriers between these states (Fig. 5, 7). This likely enables frequent transitions and suggests that they continuously interrogate membrane-based substrates, in line with their established roles in signalling and quality control (*10, 24–29*). In contrast, PARL occupies a markedly open conformational ensemble, with minimal difference between apo and substrate-bound states (Fig. S9). This behaviour points to a mechanism in which substrate cleavage is less regulated by lateral gating, but is instead predominantly regulated by subcellular colocalisation of rhomboid and substrate. Such a mechanism is consistent with the role of PARL in the constitutive processing of PINK1 in the IMM of healthy mitochondria (*22, 49*).

In striking contrast, the orphans RHBDL1 and RHBDL3 exhibit narrower lateral gates, rarely occupying states consistent with lateral gating (Fig. 6-7). Both equilibrium and enhanced sampling simulations show that open states are energetically disfavoured, despite being structurally accessible. This places the orphan rhomboids in a distinct category in which substrate access is constrained. A key advance of this work is the explicit linkage between lateral gate width and the associated free energy landscape. While all rhomboids are capable of adopting conformations compatible with substrate entry (∼ 1.4 nm or greater), the likelihood of sampling these states differs substantially. RHBDL4 and PARL energetically favour open conformations; RHBDL2 displays multiple minima separated by modest energy barriers; and the orphans are dominated by deep energy minima corresponding to closed states (Fig. 7). We therefore propose that substrate access is best described as a probabilistic process governed by how frequently a given rhomboid protease samples a gating-competent conformation, reframing lateral gating as an energy landscape problem, rather than a purely structural one.

Structural features observed in the models provide potential explanations for these differences. Most notably, RHBDL4 and PARL share unique features not seen in the other rhomboids, perhaps reflecting a shared molecular function. These include shorter TM helices, a broken TM6 and a beta-sheet in the loop between TM4 and TM5, which sits in a hotspot for dynamics in RHBDL4/PARL MD simulations (Fig. S7, S9G, S15). These features may stabilise the lateral gate in more open conformations and reduce the energetic cost of gate opening. This would fit with their quality control roles at sites of membrane protein insertion (the ER and inner mitochondrial membrane, respectively), where they survey their environment for misfolded or mislocalised proteins(*27, 30*). More broadly, variation in transmembrane helix length likely reflects a known adaptation of membrane proteins to differential membrane thickness in subcellular compartments(*53*), suggesting that membrane environment and protein structure co-evolve to tune gating behaviour.

Together, these findings support a revised model of rhomboid proteolysis in which lateral gate opening occurs stochastically, and substrates opportunistically sample the rhomboid fold. Productive cleavage likely arises from the temporal alignment of gate opening with substrate helix destabilisation, rather than from a strictly substrate-induced conformational change in the rhomboid fold. In this framework, proteolysis emerges from the coincidence of enzyme and substrate dynamics.

The constrained conformational landscapes of RHBDL1 and RHBDL3 provide a compelling explanation for their status as orphan proteases, yet they also suggest that activity may be conditionally enabled through mechanisms that transiently overcome these structural restrictions. First, their high energetic barriers to lateral gate opening suggest that substrate access may require additional stabilising interactions or regulatory inputs. The absence of a second energy minimum at ∼1.2 nm (as seen for RHBDL2; Fig. 7E) in the energy landscape of orphan rhomboids may point to an unprecedented requirement of a lateral gating cofactor for these rhomboid proteases. Such a cofactor may lower the energetic barrier to gate opening, effectively priming the fold for substrate engagement. Although all characterised rhomboid proteases to date are capable of cleaving substrates without cofactors, this should not be assumed for the orphans. The plausibility of a membrane-bound rhomboid regulator is supported by the observation that rhomboid family members, including pseudoproteases, can form functional units with other membrane proteins (*49, 54, 55*).

Second, the heterogeneity in cytoplasmic domains across the rhomboid superfamily indicates distinct modes of regulation for localisation, interaction partners and conformational dynamics, which are all likely related to their overall roles. We found that all rhomboids modelled using MD featured unusually dynamic intracellular domains that remained highly flexible throughout the entire simulation runs (Figure S8). Therefore, a key factor to consider is the extended cytoplasmic domains of the orphans, which may function allosterically, coupling distal signals to conformational transitions within the membrane. Both RHBDL1 and RHBDL3 share orphan-specific extended N-terminal domain containing EF hands domains (Fig. 2, Fig. S16) (*56*). Stabilisation or regulation of the orphan cytoplasmic domains could propagate conformational changes and lock the proteases into either an open or closed state. This is notably the case for Drosophila rho-4, which requires calcium regulation of cytoplasmic regions for protease activity (*57*).

Third, it is possible that RHBDL1 and RHBDL3 process a distinct class of protein to that of canonical rhomboid substrates, namely single-pass type I membrane proteins. A substrate which can engage the active site without full opening of the lateral gate would fit this profile, similar to the processing of soluble substrates by RHBDL4 or GlpG (*30, 58*). Interestingly, RHBDL1 and RHBDL3 diverged from RHBDL2 relatively late in evolution, raising the possibility that these proteases are adapted to cleave proteins that specifically emerged in metazoans. Given that the evolution of a nervous system is a defining feature of metazoan biology, RHBDL1 and RHBDL3 may have specialised roles in processing neuronal proteins, consistent with their endogenous expression patterns (*33, 34, 59*). For these reasons, substrate identification efforts for RHBDL1 and RHBDL3 should be focussed on these cell types.

More broadly, this work underscores the importance of integrating AI-based structural prediction with physics-based simulations to capture a full picture of conformational ensembles, particularly for membrane proteins. While static models provide important structural insights, it is the distribution of accessible states and their energetic relationships that ultimately define their mechanism of action. The rhomboid proteases provide a clear example of how tuning of conformational energy landscapes can regulate biological functions, a principle that likely extends to a broad variety of polytopic proteins such as GPCRs, ion channels and transporters.

## Methods

### Structural prediction

AlphaFold models were generated using AlphaFold2 ColabFold, using a PDB template and relaxation(*60*). Chai-1 models were generated using the webserver, with an MMseqs2 MSA and templates(*37*). Boltz-2 predictions were performed locally using default settings. Coordinate files and confidence scores can be found at https://osf.io/n6348/. For PARL:PINK1, the models reported previously were used(*49*). Sequences for used for all models are in Table 1. Images were generated using ChimeraX(*61*). Structural alignments and RMSD values for core fold comparison between models generated by different tools were performed using ChimeraX using the Matchmaker tool. The residues used to define the core fold for these purposes are listed in Table 2.

**Table 1-.** Input sequences used to generate AF2 models.

| Predictive model | Input sequence |
| --- | --- |
| GlpG | All residues of UniProt entry P09391 |
| RHBDL1 | All residues of UniProt entry O75783-2 |
| RHBDL2 | All residues of UniProt entry Q9NX52 |
| RHBDL3 | All residues of UniProt entry P58872 |
| RHBDL4 | All residues of UniProt entry Q8TEB9 |
| PARL | Residues 78-379 of UniProt entry Q9H300 |
| RHBDL2:Orai | All residues of UniProt entry Q9NX52;<br>residues 233-257 of Q96D31 |
| RHBDL4:pTa | All residues of UniProt entry Q8TEB9;<br>residues 145-169 of Q6ISU1 |
| PARL:PINK1 | Residues 93-368 of UniProt entry Q9H300;<br>all residues Q9BXM7 |

**Table 2-.**
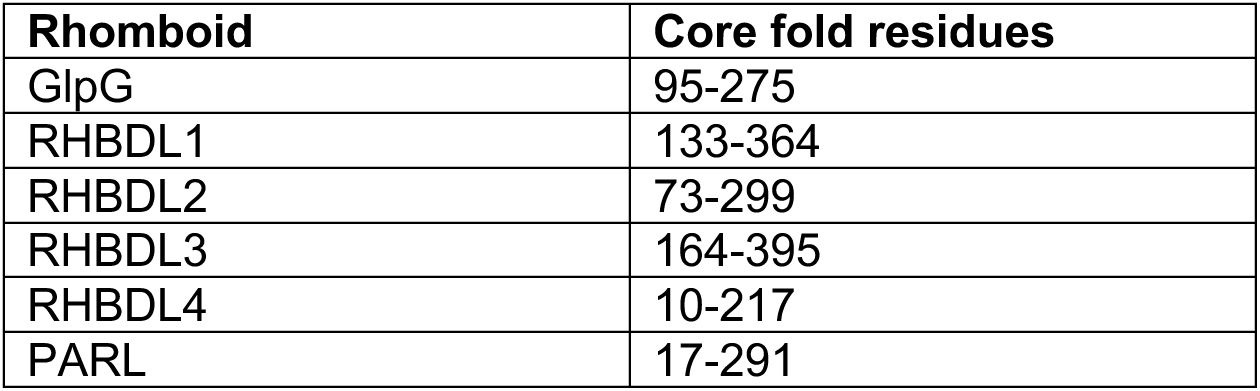
Residues used to calculate RMSD values between rhomboid models from different structure prediction tools.

| Rhomboid | Core fold residues |
| --- | --- |
| GlpG | 95-275 |
| RHBDL1 | 133-364 |
| RHBDL2 | 73-299 |
| RHBDL3 | 164-395 |
| RHBDL4 | 10-217 |
| PARL | 17-291 |

### Lateral gate quantification

To systematically compare lateral gate distances across AF2 models, rhomboids were structurally aligned in ChimeraX and 2 residue pairs which were positioned similarly along transmembrane helices 2 and 5, with one pair above and one pair below the active site. The distance between the alpha-carbon atoms was measured using the measurement wizard and the mean was calculated. Data were then plotted using Prism 10. Residue pairs were as displayed in Table 3.

**Table 3-.** residue pairs used for lateral gate quantification of rhomboid models/structures. Equivalent residues were also used for quantification of MD simulations, using Gromacs 2024.4 (62).

| Rhomboid | Residue pair 1 | Residue pair 2 |
| --- | --- | --- |
| GlpG | G240-M149 | W236-N154 |
| RHBDL1 | R290-E194 | E287-N199 |
| RHBDL2 | F230-Q134 | D227-N139 |
| RHBDL3 | R321-E225 | E318-N230 |
| RHBDL4 | I183-W79 | E179-N84 |
| PARL | D241-F142 | I237-Q147 |

### MD simulations

The top-ranking AF2 model was used for MD simulations and generated according with Table 4. CHARMM-GUI Membrane Builder (*63*) was used to generate the simulation system under a CHARMM36m force field(*64*). Membranes were built from a 3:7 cholesterol:POPC lipid composition using the default parameters. Systems underwent minimisation and equilibration, using the standard CHARMM-GUI protocol (*65, 66*). Simulations were run using Gromacs 2024.4 (*62*) and performed using the NPT ensemble with V-rescale temperature and Parrinello-Rahman pressure coupling parameters respectively and a timestep of 2 fs. 5 repeats of 2 µs were performed for equilibrium GlpG and RHBDL1-3 simulations. For all other simulations, 5 repeats of 500 ns were run.

**Table 4-.** input models used to seed MD simulations.

| MD model | Input model |
| --- | --- |
| GlpG | GlpG AF2 model |
| RHBDL1 | RHBDL1 AF2 model |
| RHBDL2 | RHBDL2 AF2 model |
| RHBDL3 | Residues 35-404 of RHBDL3 AF2 model |
| RHBDL4 | RHBDL4 AF2 model |
| RHBDL2:Orai | Only the RHBDL2 atoms from the RHBDL2:Orai AF2 model |
| RHBDL4:pTa | Only the RHBDL4 atoms from the RHBDL2:Orai AF2 model |
| PARL | Residues 16-292 of PARL AF2 model (94-370 of Q9H300) |

Lateral gate helix distance profiles were generated using the earlier described lateral gate quantification. A time step of 2.5 ns was used and a 2^nd^ order smoothing with 4 neighbours was used for the individual run traces. Histograms were generated by appending the distances across 5 simulations into one column and performing a frequency distribution analysis, using Prism. The centre of each bin was plotted along the x axis, and frequency using the 2.5 ns timestep along the y-axis. Violin plots were also generated from the appended tables, and a ‘high’ smoothing setting was applied.

### Validation of models

QMEANBrane was used to assess the validity of AF2 models, using the Expasy webserver(*40*). Sequences in Table 1 were used as inputs. MolProbity was also used to assess the quality of MD input models listed in Table 4, in addition to the 2IC8 and 2NRF GlpG structures for comparison. EVcouplings was used to analyse the co-evolution of rhomboids(*42*). The sequences listed in Table 1 were used as input queries. Rank 1 AF2 models were then used for 3D mapping of coupling scores.

### Principal component analysis

AF2 models of human RHBDL1, RHBDL2 and RHBDL3 were aligned to identify atoms conserved atoms across the core fold, which are shown in Table 5. These residues were taken forward to process trajectories of AF2 MD simulations into one concatenated simulation, with only the conserved C-alpha atoms from these trajectory files used for PCA. Using Gromacs tools, PCA was performed to identify the largest scale movements seen across the first 500 ns simulation of equilibrium MD simulations, across all 5 repeats for each rhomboid. First, a covariance matrix was built and diagonalised before the eigenvectors were projected onto each of the rhomboid simulations, allowing them to be plotted against their eigenvalues. For eigenvector 1, the two extreme states represented by this eigenvector were overlaid.

**Table 5-.** Residues used in PCA for RHBDL1-3.

| AF2 model | Residues used in PCA |
| --- | --- |
| RHBDL1 | 100-110, 112-299, 302-365 |
| RHBDL2 | 41-303 |
| RHBDL3 | 131-141, 143-330, 333-396 |

### Steered MD

Minimised and equilibrated systems of RHBDL1-4 were subjected to steered MD using Gromacs, as above, using Nose-Hoover and Parrinello-Rahman temperature and pressure coupling. A collective variable (CV) was defined as 3 alpha-carbon atoms on each of TM2 and TM5, as in Table 6. A constant velocity of 35 nm/ms was applied in the x and y coordinate axes, and simulations were run for 325 ns (RHBDL2); 400 ns (RHBDL1 and RHBDL3); or 260 ns (RHBDL4). 3 repeats were run and assessed visually in VMD(*67*) to check for artefacts such as bending of helices. RMSD and x-coordinate plots were also assessed to check for smooth changes throughout pulling simulations. The best of the 3 against these metrics was then taken forward for umbrella sampling simulations.

**Table 6-.** Residues used as collective variable for steered MD.

| Rhomboid | TM2 | TM5 |
| --- | --- | --- |
| RHBDL1 | E194, F198, L202 | E287, R290, L294 |
| RHBDL2 | Q134, G138, L146 | I219, L226, F230 |
| RHBDL3 | Q225, L229, and L233. | E318, R321, L325. |
| RHBDL4 | W79, F83, S87 | E179, A182, F186 |

### Potential of mean force calculations

Following steered MD to force apart TM2 and TM5, umbrella windows were selected, with ∼0.05 nm spacing, using the pmf.py script written by Dr. Owen Vickery (DOI: 10.5281/zenodo.3592318). This produced 29 windows for RHBDL1; 24 for RHBDL2; 30 windows for RHDBL3; and 18 for RHBDL4. Each window was equilibrated for 10 ps, followed by a production run of 50 ns. An additional 50 ns was run to check for convergence and, in all instances, the original 50 ns showed good convergence and was therefore used for subsequent analyses.

The same script was then used to perform potential of mean force (PMF) calculations. The weighted histogram analysis method (WHAM) was then applied to generate an energy landscape (*50*).

## Acknowledgments

We thank Mark Dodding (University of Bristol) for proof-reading the manuscript and providing critical feedback. We also thank Philip Hinchcliffe (University of Bristol) for useful conversations. BRJC is funded via the Wellcome Trust Dynamic Molecular Cell Biology PhD programme (218510/Z/19/Z). AGG is funded by a Sir Henry Dale fellowship jointly funded by the Wellcome Trust and Royal Society (221784/Z/20/Z). This project made use of time on ARCHER2 granted via the UK High-End Computing Consortium for Biomolecular Simulation, HECBioSim (http://hecbiosim.ac.uk), supported by EPSRC (EP/R029407/1). The authors acknowledge the use of resources provided by the Isambard 3 Tier-2 HPC Facility. Isambard 3 is hosted by the University of Bristol and operated by the GW4 Alliance (https://gw4.ac.uk) and is funded by UK Research and Innovation; and the Engineering and Physical Sciences Research Council (EP/X039137/1).

## Author contributions

Conceptualisation: BRJC, RAC, AGG Investigation: BRJC Methodology: BRJC, RAC Supervision: RAC, AGG Visualisation: BRJC, RAC, AGG Writing: BRJC, RAC, AGG

## Data and materials availability

PDB files for all AF2, Boltz-2 and Chai-1 models, including confidence scores, can be found at the following online repository: https://osf.io/n6348/.

All other data are available in the main text or the supplementary materials.

