## Supplementary Materials for "Structural and energetic insights into human rhomboid proteases reveal a unique lateral gating mechanism for orphan family members"

#### Supplementary Figure Legends

**Fig. S1 – Lateral gate quantification of GlpG simulations.** Individual traces for the five MD runs shown in Fig. 1 are depicted. Dashed lines indicate the lateral gate widths of PDB entries 2NRF or 2IC8; or the rank 1 AF2 model as labelled.

**Fig. S2 – RMSD and RMSF analysis of GlpG simulations.** (Top) RMSD analysis of 5 x 2 μs equilibrium MD simulations for each GlpG structure/model. Each simulation is shown as a separate line. Analyses are with vs without intracellular domains as indicated. (Bottom) RMSF analysis of GlpG simulations. Plots show RMSF values for the alpha-carbon of each residue as an average across the 5 repeats for the entire simulation length. RMSF values are then projected onto the cartoon representation of the input model.

**Fig. S3 – Three independent structure prediction tools generate similar models for each of the human rhomboids.** Structural alignments of the highest ranked model generated by AF2, Chai-1 and Boltz-2. Only the core rhomboid fold was used for the alignment, and RMSD values against the AF2 model were calculated.

**Fig. S4 – Confidence scores for AF2 predictions. (A)** pTM scrores for the top 5 ranks of AF2 models for human rhomboids and *E. coli* GlpG. Line = mean. **(B-G)** AF2 pLDDT heatmaps for all rhomboids modelled. **(H)** Lengths of each TM of AF2 models for human rhomboids, measured from the alpha-carbon atom of the first and last helical residue (as determined by visual analysis of AF2 predictions in ChimeraX). Line = mean.

**Fig. S5 – Evolutionary coupling analysis validates the quality of AF2 models.** For each of the AF2 models used for MD simulations, EVcouplings was used to project the cumulative coupling strength per site onto the rank 1 AF2 model(*42*). The scale bar for each heat map is shown below the model, with blue indicating residues with higher cumulative coupling scores.

**Fig. S6 – Evolutionary coupling analysis validates the quality of AF2 models.** For each of the AF2 models used for MD simulations, EVcouplings was used to generate contact maps, with green circles showing points of contact within the 3D model. Black dots indicate an evolutionary coupling and are scaled to the strength of this coupling.

**Fig. S7 – RHBDL4 and PARL share similar TM features** Cartoon depictions of rank 1 AF2 models of RHBDL4 (green) and PARL (brown). **(A)** Overlay of RHBDL4 and PARL. Black arrow indicates break in TM6. **(B,C)** Cartoon depictions of RHDBL4 and PARL respectively, as for (A) but showing the cholesterol and POPC atoms from the CHARMM-GUI generated lipid bilayer (7:3 POPC:cholesterol composition), following equilibration. **(D)** Overlay of RHBDL4 and PARL. View is looking into the lateral gate. Positions of lateral gate helices are indicated. Black box shows a zoomed in view of the shared beta-sheet feature.

**Fig. S8 – RMSD plots from MD simulations.** RMSD analysis equilibrium MD simulations for each rank 1 RHBDL1-4 AF2 model as indicated. Each simulation is shown as a separate line. Analyses are with or without intracellular domains as indicated.

**Fig. S9 – Molecular dynamics simulations of PARL. (A)** Lateral gate width profiles for human PARL during molecular dynamics simulations. The rank 1 AF2 model was used as an input following generation of a 7:3 POPC:cholesterol lipid bilayer and energy minimisation. Five 500 ns simulations were performed. Each simulation is shown as a separate line. Dashed lines show the input lateral gate width. **(B,C)** Histograms showing the distribution of lateral gate widths across equilibrium simulations of PARL compared to RHBDL4 and RHBDL2 respectively. **(D)** Violin plots showing the distribution of lateral gate widths across equilibrium simulations of PARL compared to RHBDL2 and RHBDL4. **(E)** RMSD plot from simulations shown in (A). **(F)** RMSF plot for backbone atoms of PARL in simulations shown in (A). **(G)** RMSF heatmap of PARL.

**Fig. S10 – Molecular dynamics simulations of RHBDL4 using an ER-like membrane composition.** Equilibrium simulations for the rank 1 RHBDL4 model (Fig. 4) were repeated using a representative ER membrane composition (45:45:10 DMPC:DMPE:cholesterol (CHOL)). **(A)** Lateral gate helix distance profiles for RHBDL4 during molecular dynamics simulations. Five 500 ns simulations were performed. Each simulation is shown as a separate line. Dashed line shows the input lateral gate width (post equilibration in a lipid bilayer). **(B)** Histograms show the lateral gate width distributions across simulations shown in (A) and Fig. 4B (ER vs plasma membrane composition). **(C)** Violin plots comparing RHBDL4 lateral gate width profiles across the two different membrane compositions**. (D)** RMSD plots for the core rhomboid fold of RHBDL4 in simulations shown in (A). **(E)** RMSD plots for the full-length RHBDL4 in a 45:45:10 DMPC:DMPE:CHOL membrane. **(F)** RMSF plot for backbone atoms of RHBDL4 in simulations shown in (A).

**Fig. S11 – Principal component analysis of RHBDL1-3 MD simulations.** Equivalent backbone atoms were identified within the core rhomboid fold across RHBDL1-3. These were selected for PCA across the first 500 ns of all equilibrium MD simulations of rank 1 AF2 models. **(A)** Eigenvalues for the first 12 eigenvectors of the covariance matrix from RHBDL1-3 simulations. **(B)** Overlay of the 2 extreme states from eigenvector 1 projected onto the RHBDL1/2/3 skeleton. Lateral gate distances are indicated. **(C)** Projections of eigenvector 1 individually onto MD trajectories of RHBDL1, -2 and -3.

**Fig. S12 – Equilibrium MD simulations of RHBDL1/3 using steered MD outputs. (A)** Distance plot showing the CV distance of RHBDL3 over the duration of the steered MD run (see Fig. 7). **(B-C)** Lateral gate width profiles for RHBDL1 and RHBDL3 respectively during molecular dynamics simulations. The final frame of steered MD was used as an input following generation of a 7:3 POPC:cholesterol lipid bilayer and energy minimisation. Five 500 ns simulations were performed. Each simulation is shown as a separate line. Dashed lines show the input lateral gate width ('post-pull'); the widest state seen in equilibrium MD simulations seeded from the AF2 rank 1 model ('Widest AF2-MD state') and the AF2 rank 1 prediction ('AF2 rank 1'). **(D)** Violin plots showing the comparison of equilibrium MD simulations seeded with AF2 models vs pulled open models.

**Fig. S13 – Histograms of umbrella sampling.** Histograms from Owen Vickory's PMF script, showing the sampling of umbrella windows for RHBDL1-4 PMF analyses.

**Fig. S14 – Molecular dynamics simulations of gated RHBDL2.** AFM predictions of gated rhomboids were seeded into simulations with atoms from the transmembrane substrate deleted. **(A)** Overlay of the structurally aligned rank1 AF2 model of RHBDL2 apo (light blue) with the AFM model of 'gated' RHBDL2 (navy blue). **(B)** Lateral gate distance profiles from five 500 ns equilibrium simulations seeded with the RHBDL2 gated model shown in (A). Each simulation is shown as a separate line. Dashed line shows the input lateral gate width (post equilibration in a lipid bilayer). **(C)** Histogram showing the frequency distribution of lateral gate widths from RHBDL2 simulations using the apo AF2 input model (Data replotted from Figure 5) or the gated AFM model. **(D)** Violin plot comparing lateral gate width distribution in simulations of the RHBDL2 apo (Data replotted from Figure 5) or gated models. The dashed line indicates the ~1.4 nm gating threshold and the cumulative frequency of simulation time above this threshold is displayed above each density curve as a percentage.

**Fig. S15 – RMSF analysis of equilibrium MD simulations of rhomboid AF models**. Plots (right) show RMSF values for full protein; heat maps (left) show analysis with intracellular domain removed (due to high flexibility, allowing flexibility within core fold to be more easily visualised). RMSF values for each residue are an average across the 5 repeats for the entire simulation length. RMSF values are then projected onto the cartoon representation of the input model.

**Fig. S16 – Predicted EF-hands domains of RHBDL1 and RHBDL3.** Due to the presence of EF-hands domains, RHBDL1 and RHBDL3 have been suggested to be regulated by calcium, as has been shown for the EF-hand containing rhomboid rho-4 in Drosophila. However, the second of these EF-hand motifs in RHBDL1 and RHBDL3 lack key calcium coordinating residues, and the general fold thus appears similar to the Eps15 homology (EH) fold. **(A-B)** N-terminal domains of RHBDL1 (orange) and RHBDL3 (purple) are shown as cartoon representations with the predicted

EF hands motif shown in red. **(C)** Overlay of structural alignment of (A) and (B). **(D)** Sequence alignments from Clustal Omega of residues 1-84 (RHBDL1) and 35-115 (RHBDL3, unstructured 1-34 region removed). RMSD = 0.31 nm. EF hands consensus motif = DxDxxGxIxxxE.

### Figure S1

#### GlpG MD lateral gate quantification

##### GlpG open (2NRF)

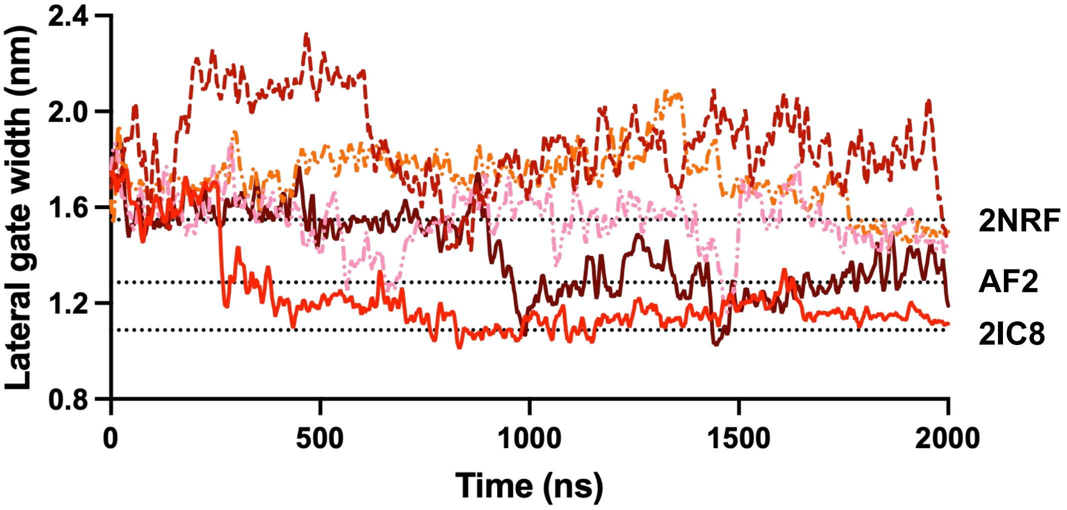


##### GlpG closed (2IC8)

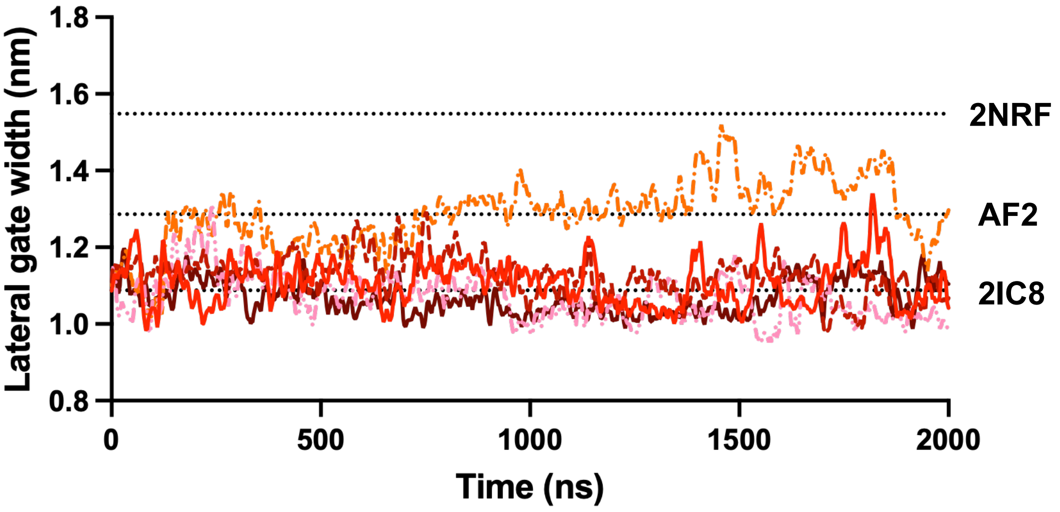


##### GlpG AF2

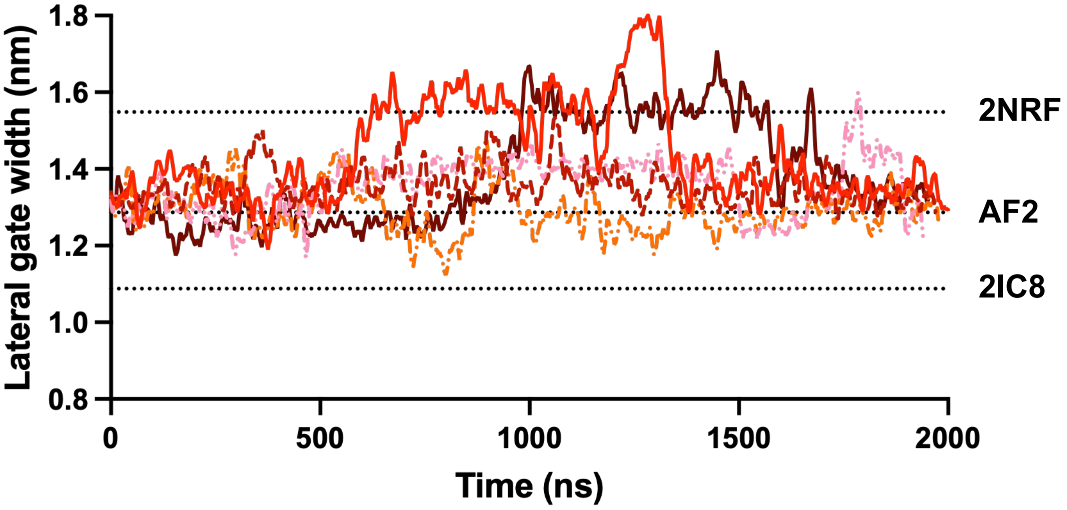

### Figure S2

#### RMSD

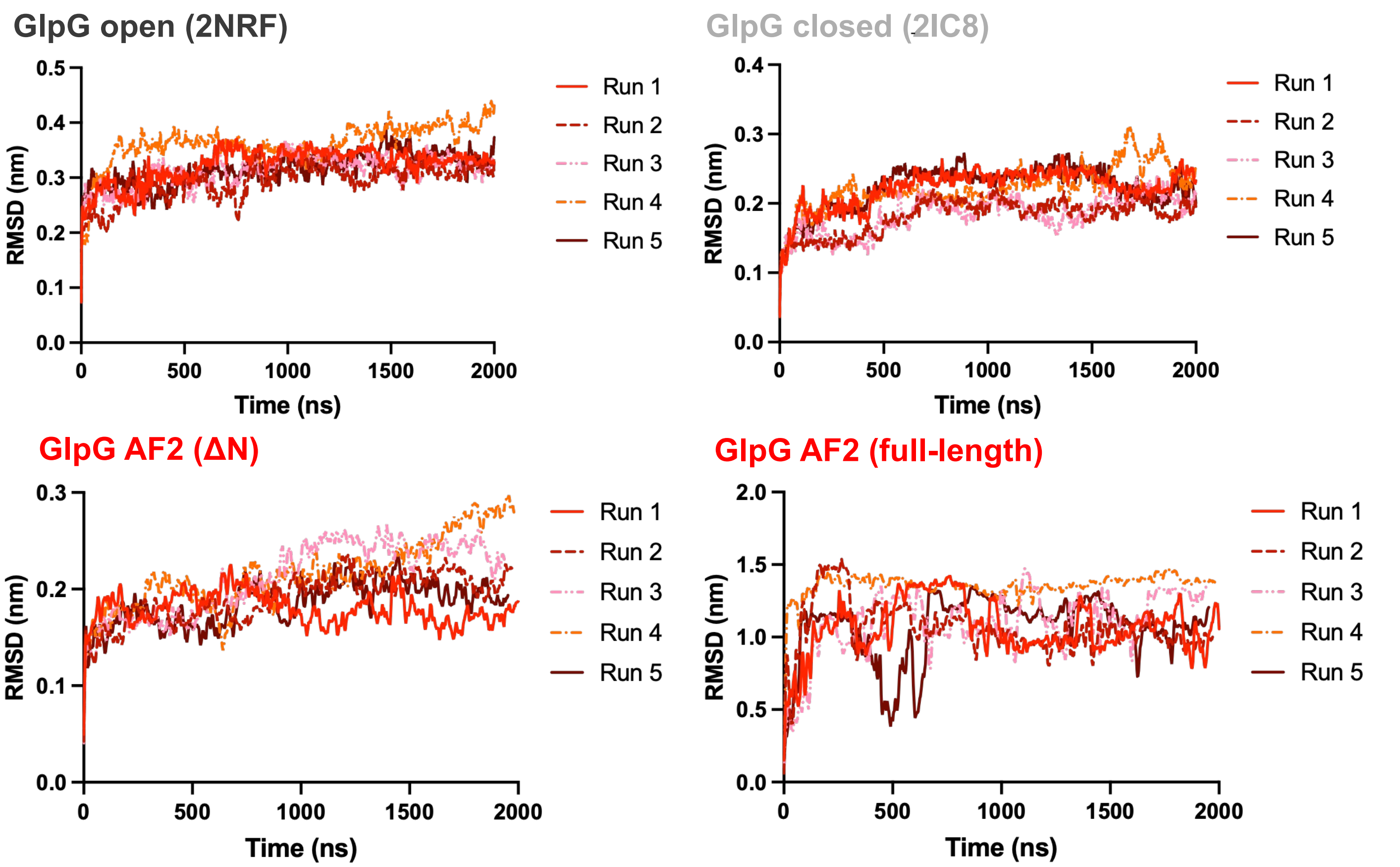


#### RMSF

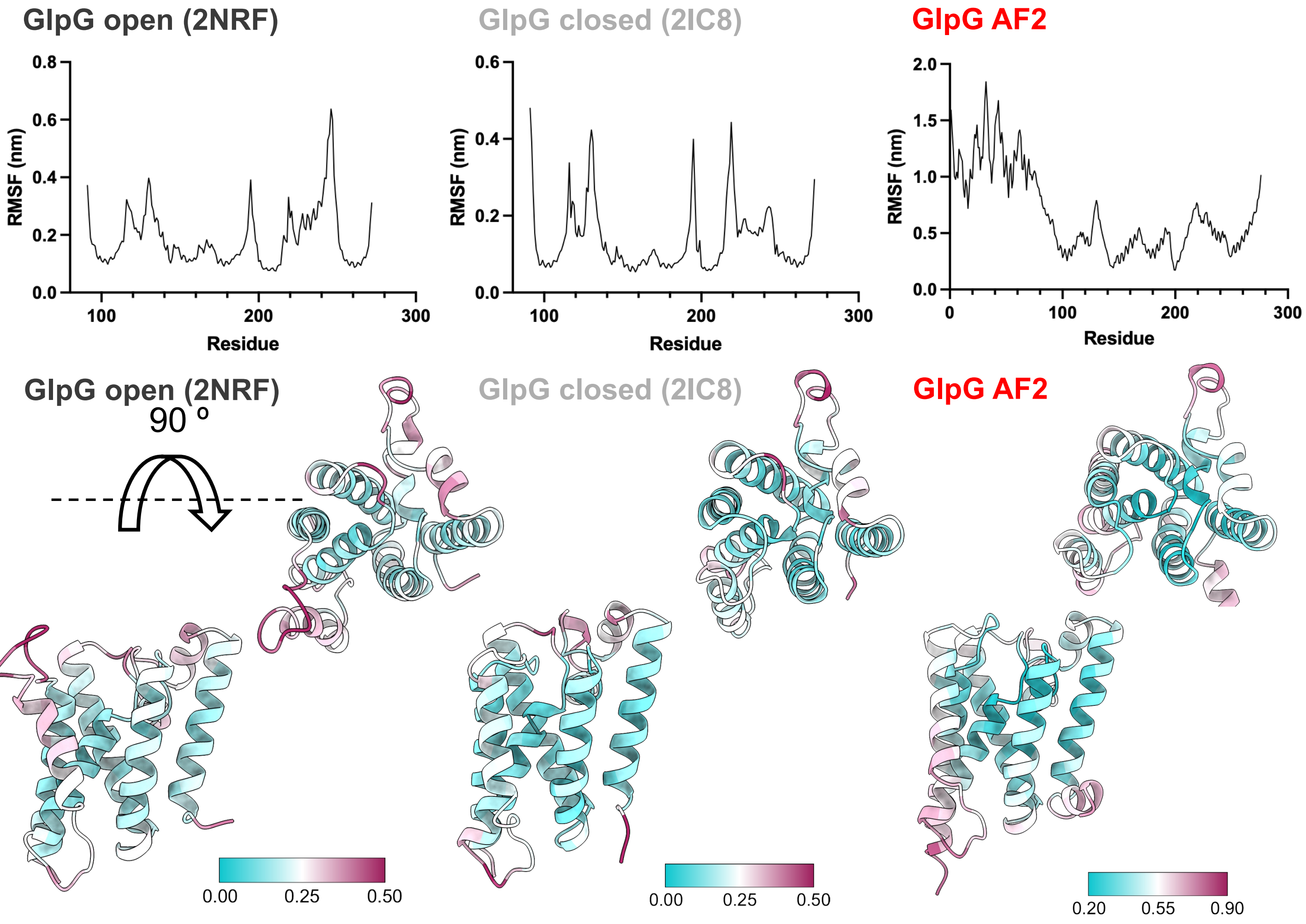

**Figure S3**

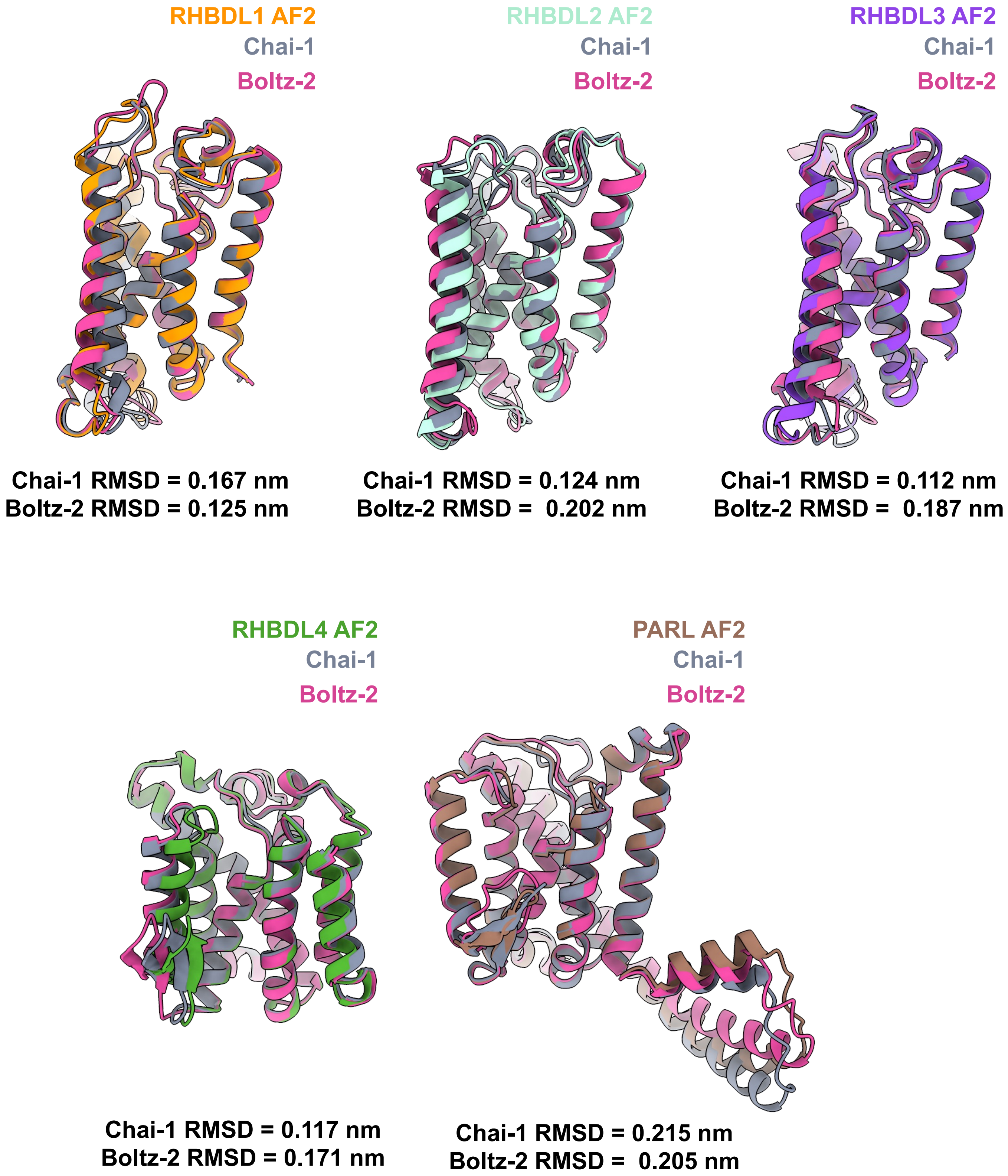
RHBDL1 AF2
Chai-1
Boltz-2
Chai-1 RMSD = 0.167 nm
Boltz-2 RMSD = 0.125 nm
RHBDL2 AF2
Chai-1
Boltz-2
Chai-1 RMSD = 0.124 nm
Boltz-2 RMSD = 0.202 nm
RHBDL3 AF2
Chai-1
Boltz-2
Chai-1 RMSD = 0.112 nm
Boltz-2 RMSD = 0.187 nm
RHBDL4 AF2
Chai-1
Boltz-2
Chai-1 RMSD = 0.117 nm
Boltz-2 RMSD = 0.171 nm
PARL AF2
Chai-1
Boltz-2
Chai-1 RMSD = 0.215 nm
Boltz-2 RMSD = 0.205 nm

### Figure S4

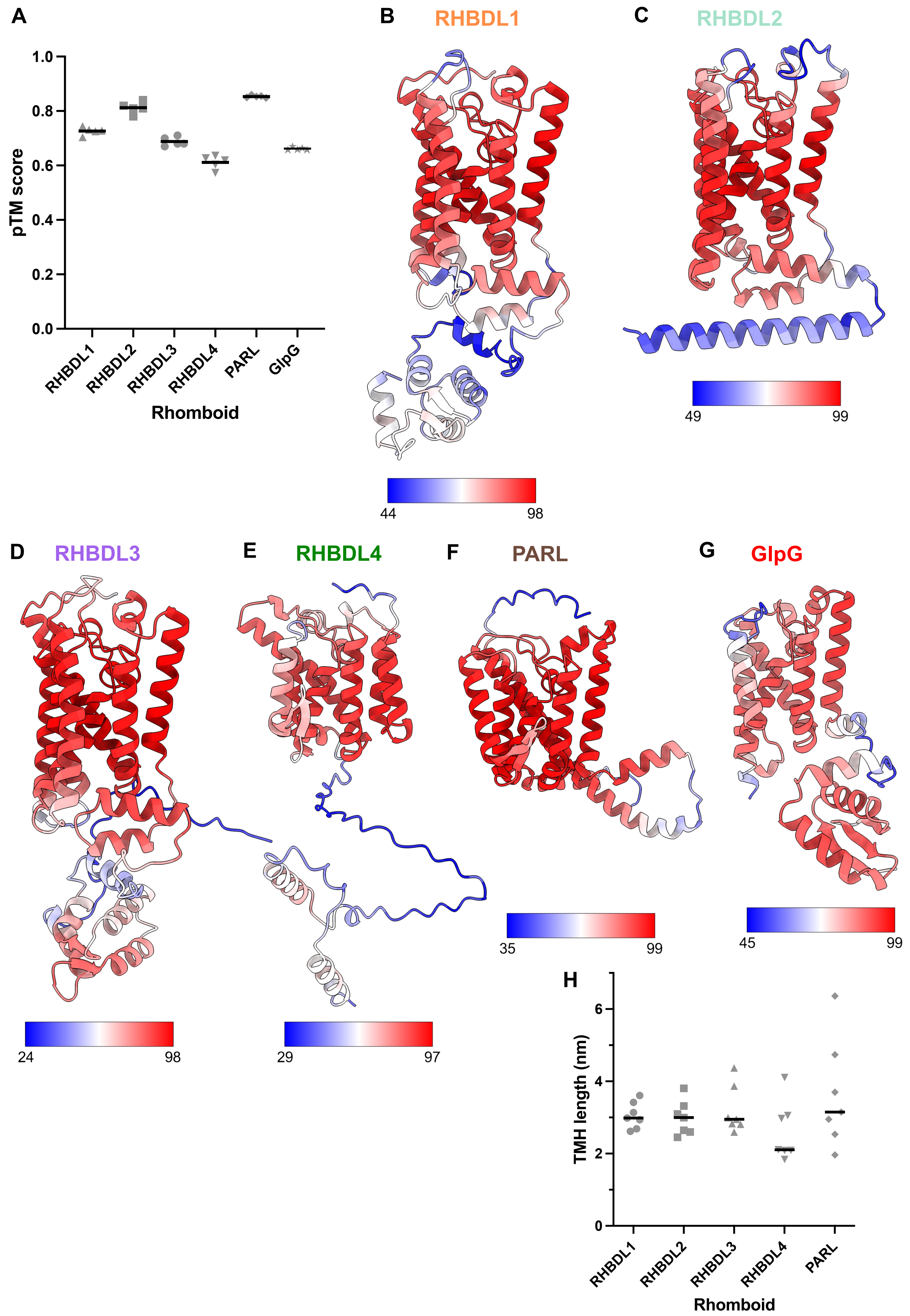

A
pTM score
1.0
0.8
0.6
0.4
0.2
0.0
RHBDL1
RHBDL2
RHBDL3
RHBDL4
PARL
GlpG
Rhomboid
B
RHBDL1
44
98
C
RHBDL2
49
99
D
RHBDL3
24
98
E
RHBDL4
29
97
F
PARL
35
99
G
GlpG
45
99
H
TMH length (nm)
6
4
2
0
RHBDL1
RHBDL2
RHBDL3
RHBDL4
PARL
Rhomboid

### Figure S5

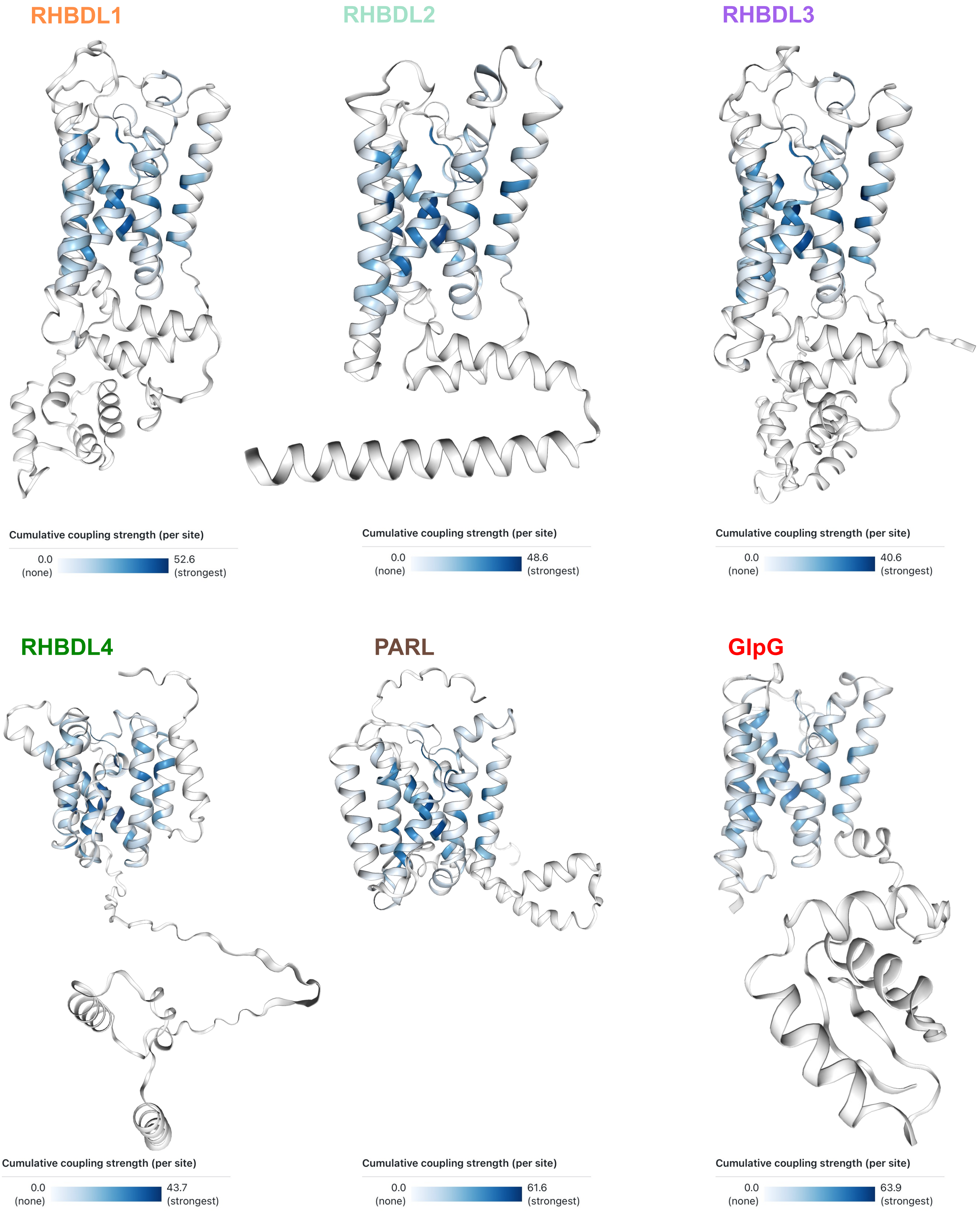
RHBDL1
Cumulative coupling strength (per site)
0.0 (none)
52.6 (strongest)
RHBDL2
Cumulative coupling strength (per site)
0.0 (none)
48.6 (strongest)
RHBDL3
Cumulative coupling strength (per site)
0.0 (none)
40.6 (strongest)
RHBDL4
Cumulative coupling strength (per site)
0.0 (none)
43.7 (strongest)
PARL
Cumulative coupling strength (per site)
0.0 (none)
61.6 (strongest)
GlpG
Cumulative coupling strength (per site)
0.0 (none)
63.9 (strongest)

### Figure S6

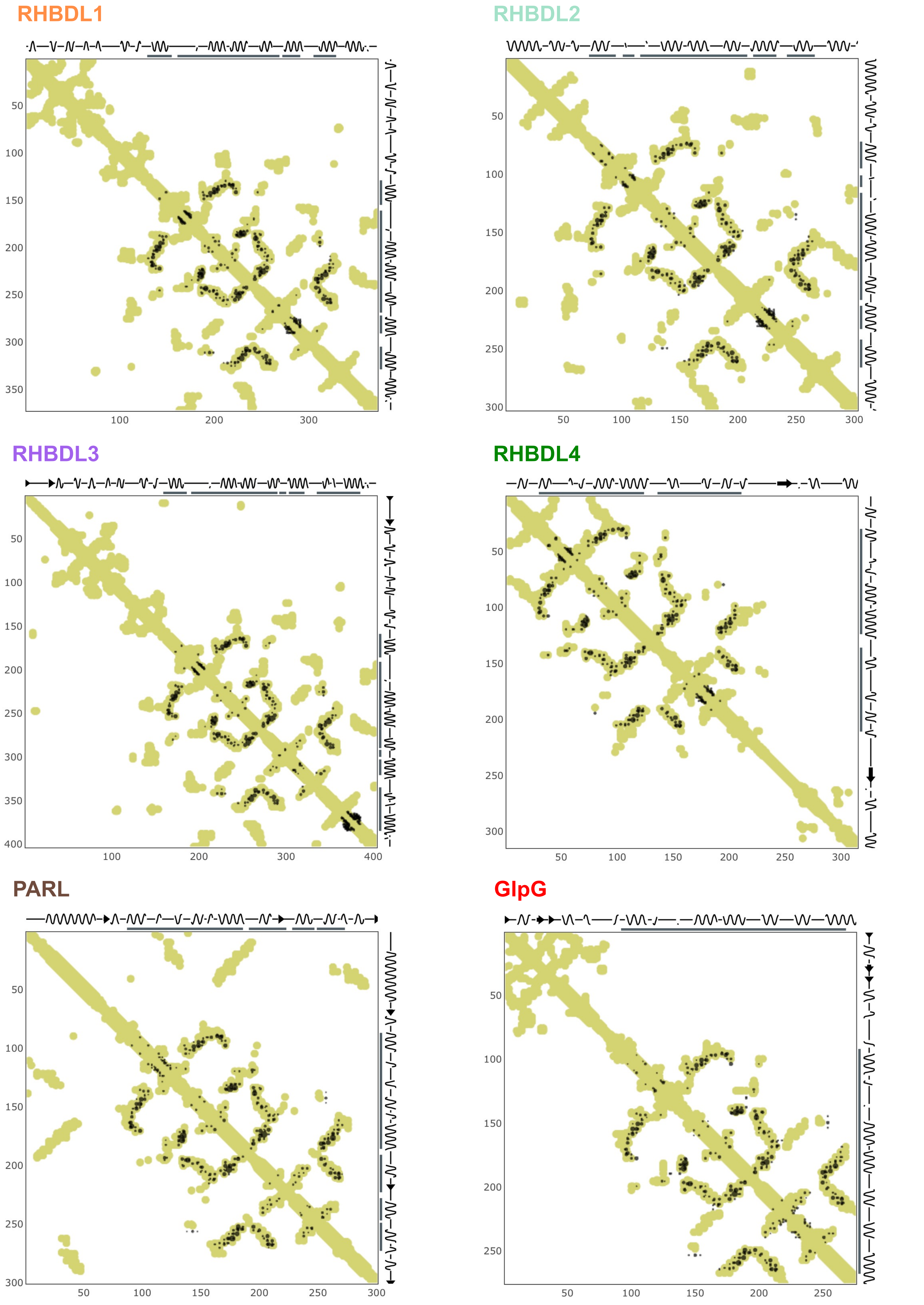

RHBDL1
50 100 150 200 250 300 350
100 200 300
RHBDL2
50 100 150 200 250 300
50 100 150 200 250 300
RHBDL3
50 100 150 200 250 300 350 400
100 200 300 400
RHBDL4
50 100 150 200 250 300
50 100 150 200 250 300
PARL
50 100 150 200 250 300
50 100 150 200 250 300
GlpG
50 100 150 200 250
50 100 150 200 250

### Figure S7

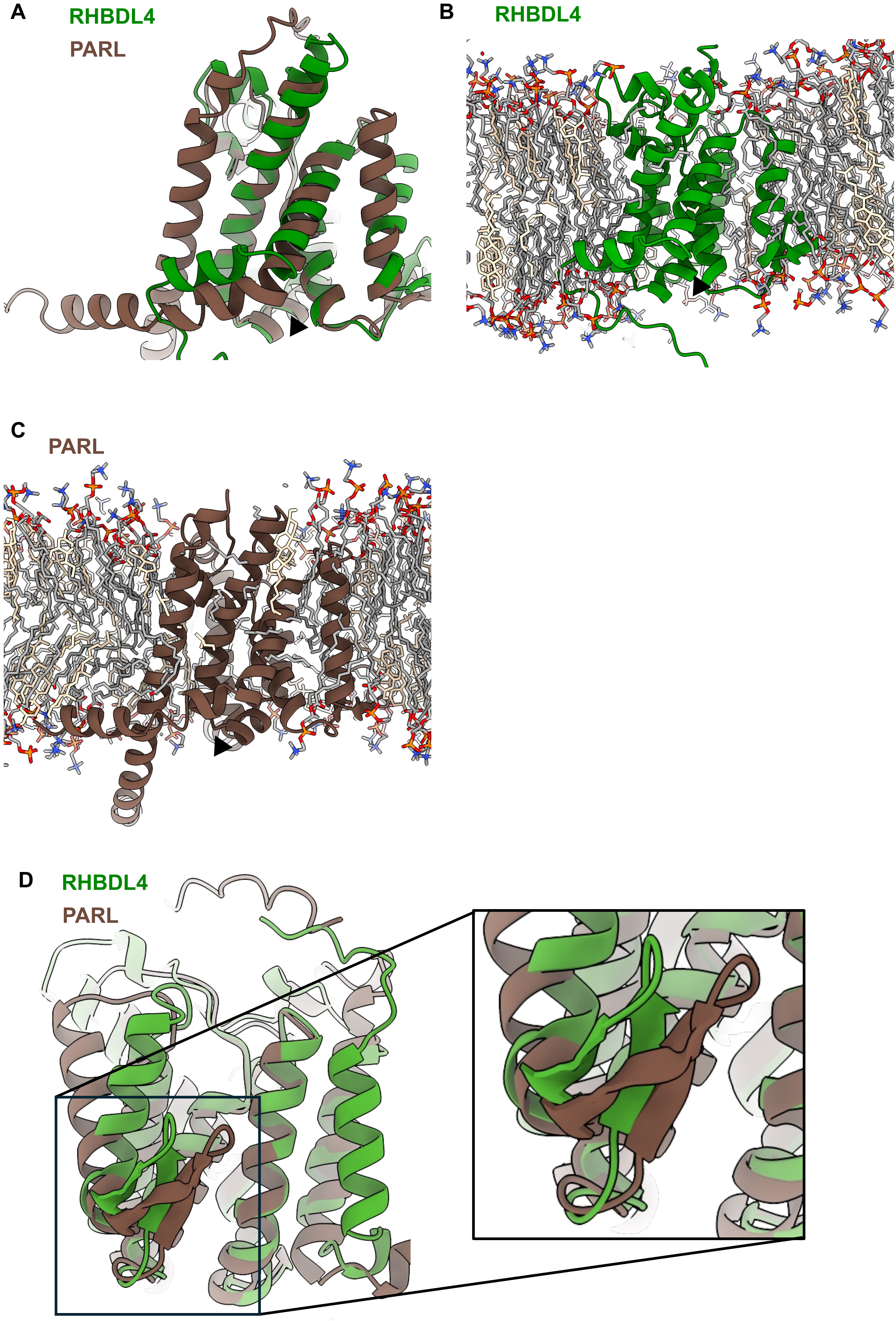

### Figure S8

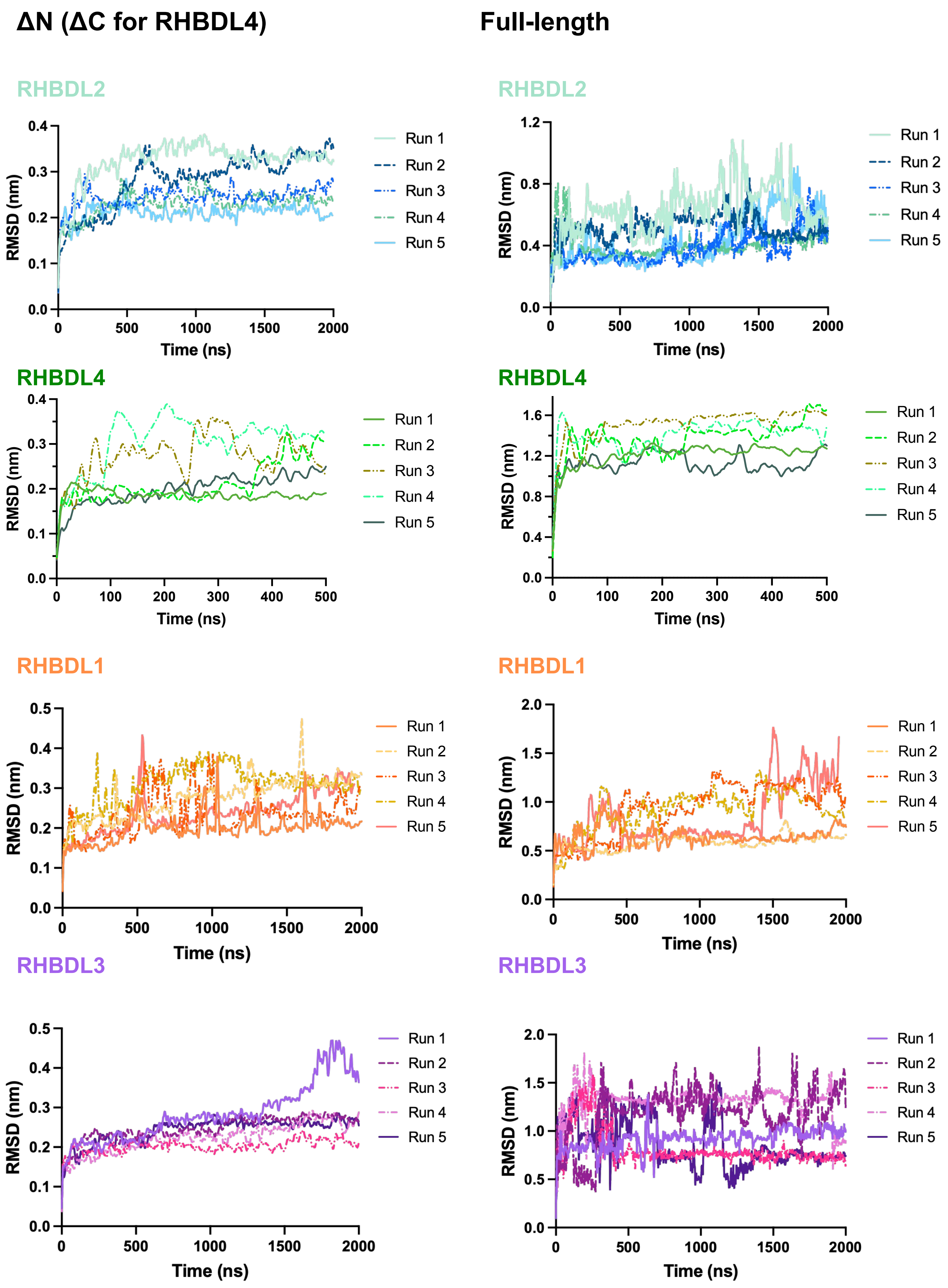
ΔN (ΔC for RHBDL4)
Full-length
RHBDL2
RHBDL4
RHBDL1
RHBDL3
RMSD (nm)
Time (ns)
Run 1
Run 2
Run 3
Run 4
Run 5
0.0
0.1
0.2
0.3
0.4
0.5
0.8
1.2
1.6
1.0
1.5
2.0
0
100
200
300
400
500
1000
1500
2000

### Figure S9

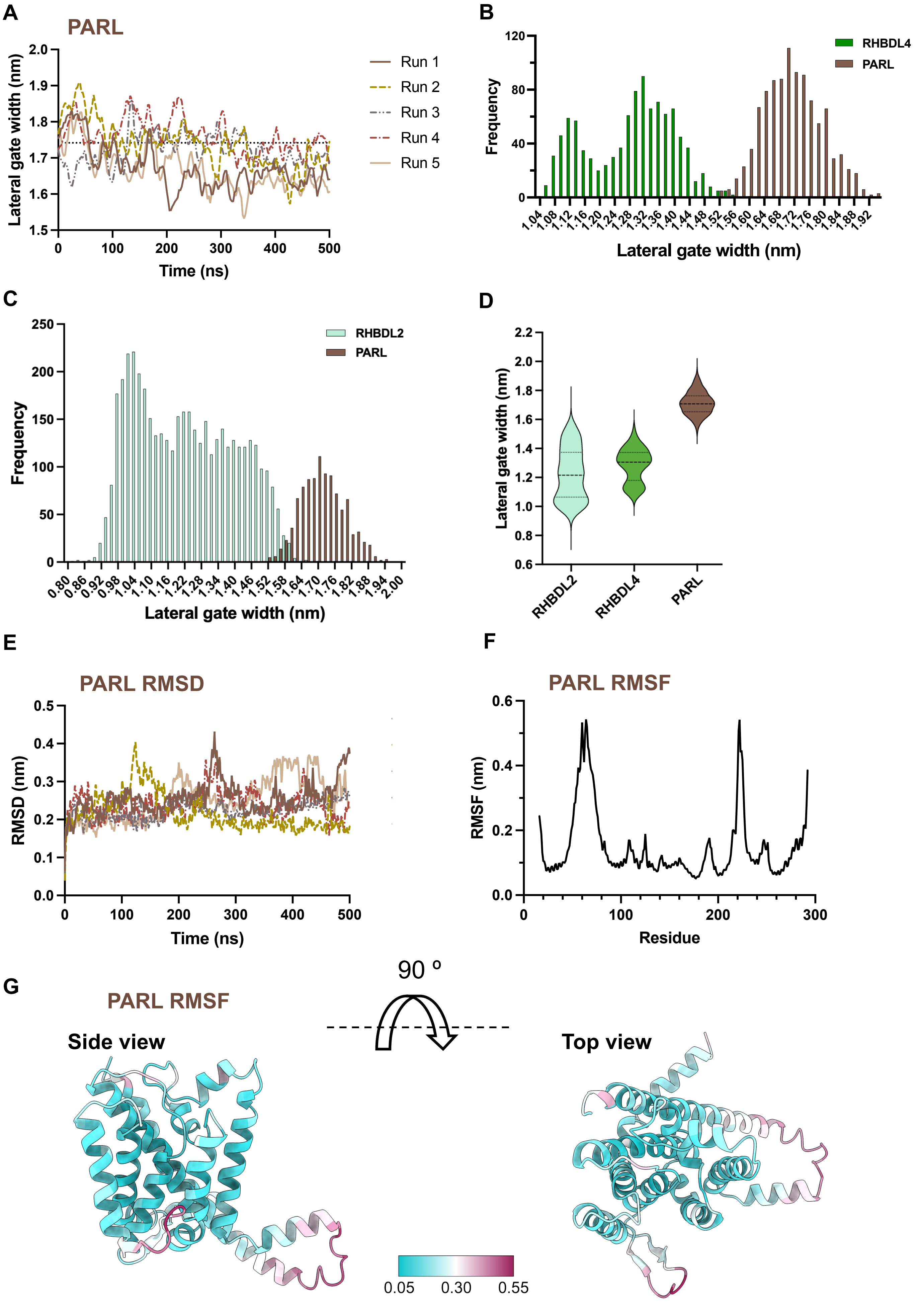
A
PARL
Lateral gate width (nm)
Time (ns)
Run 1
Run 2
Run 3
Run 4
Run 5
B
RHBDL4
PARL
Frequency
Lateral gate width (nm)
C
RHBDL2
PARL
Frequency
Lateral gate width (nm)
D
Lateral gate width (nm)
RHBDL2
RHBDL4
PARL
E
PARL RMSD
RMSD (nm)
Time (ns)
F
PARL RMSF
RMSF (nm)
Residue
G
PARL RMSF
Side view
90 º
Top view
0.05
0.30
0.55

### Figure S10

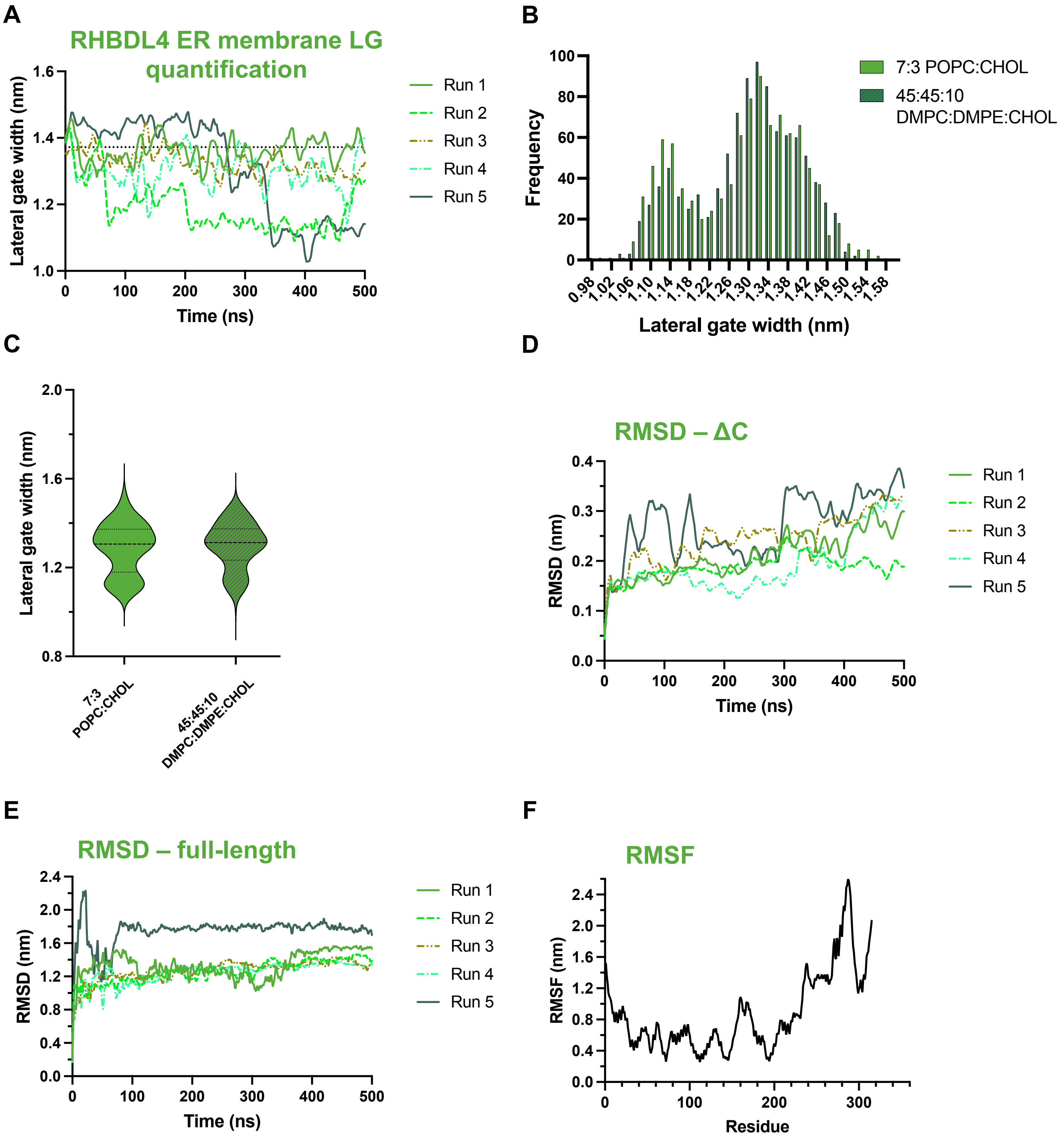

A
RHBDL4 ER membrane LG quantification
Lateral gate width (nm)
Time (ns)
Run 1
Run 2
Run 3
Run 4
Run 5
B
Frequency
Lateral gate width (nm)
7:3 POPC:CHOL
45:45:10 DMPC:DMPE:CHOL
C
Lateral gate width (nm)
7:3 POPC:CHOL
45:45:10 DMPC:DMPE:CHOL
D
RMSD – ΔC
RMSD (nm)
Time (ns)
Run 1
Run 2
Run 3
Run 4
Run 5
E
RMSD – full-length
RMSD (nm)
Time (ns)
Run 1
Run 2
Run 3
Run 4
Run 5
F
RMSF
RMSF (nm)
Residue

### Figure S11

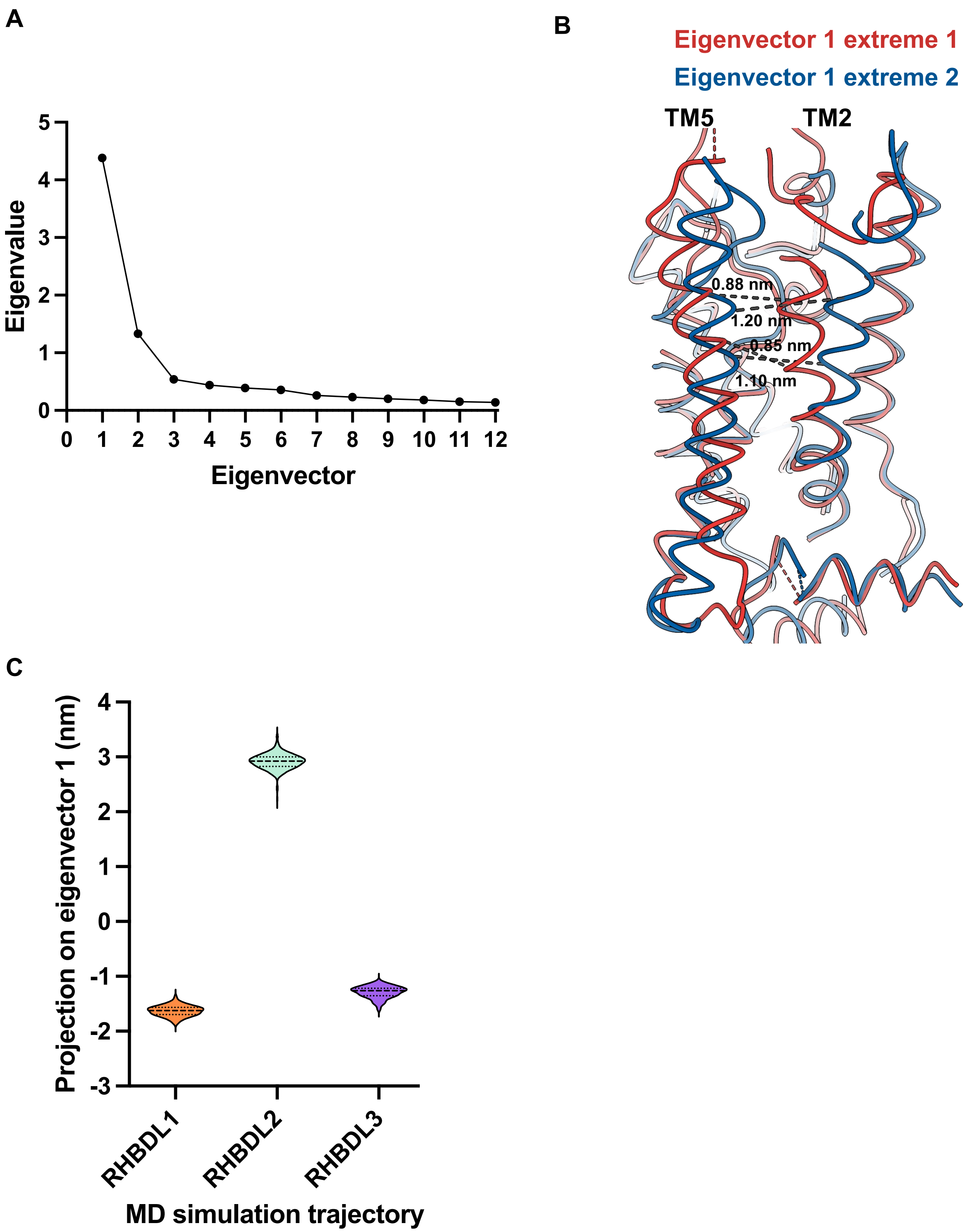

A
Eigenvalue
5
4
3
2
1
0
0 1 2 3 4 5 6 7 8 9 10 11 12
Eigenvector
B
Eigenvector 1 extreme 1
Eigenvector 1 extreme 2
TM5
TM2
0.88 nm
1.20 nm
0.85 nm
1.10 nm
C
Projection on eigenvector 1 (nm)
4
3
2
1
0
-1
-2
-3
RHBDL1
RHBDL2
RHBDL3
MD simulation trajectory

### Figure S12

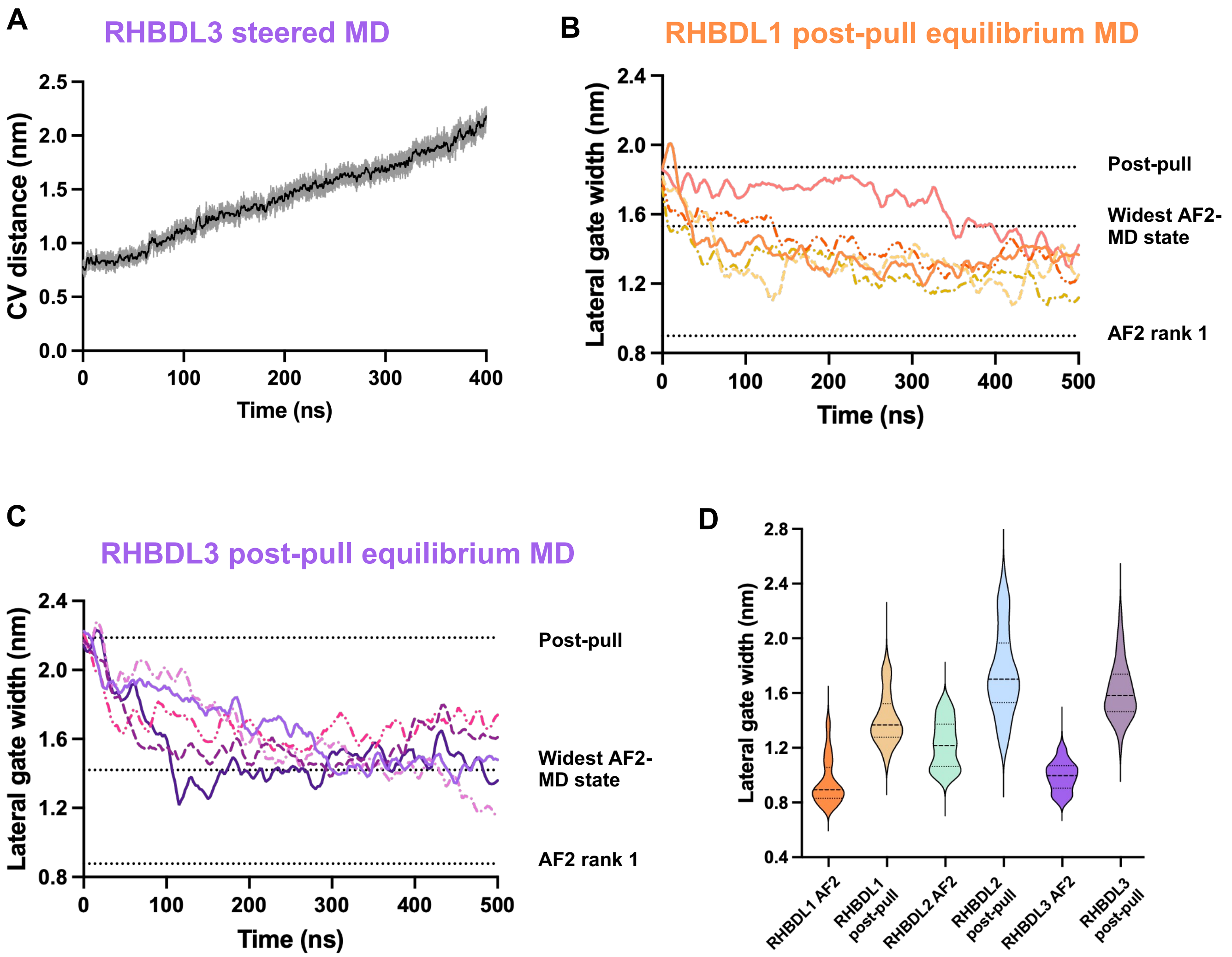

A
RHBDL3 steered MD
CV distance (nm)
2.5
2.0
1.5
1.0
0.5
0.0
0
100
200
300
400
Time (ns)
B
RHBDL1 post-pull equilibrium MD
Lateral gate width (nm)
2.4
2.0
1.6
1.2
0.8
0
100
200
300
400
500
Time (ns)
Post-pull
Widest AF2-MD state
AF2 rank 1
C
RHBDL3 post-pull equilibrium MD
Lateral gate width (nm)
2.4
2.0
1.6
1.2
0.8
0
100
200
300
400
500
Time (ns)
Post-pull
Widest AF2-MD state
AF2 rank 1
D
Lateral gate width (nm)
2.8
2.4
2.0
1.6
1.2
0.8
0.4
RHBDL1 AF2
RHBDL1 post-pull
RHBDL2 AF2
RHBDL2 post-pull
RHBDL3 AF2
RHBDL3 post-pull

### Figure S13

#### RHBDL1

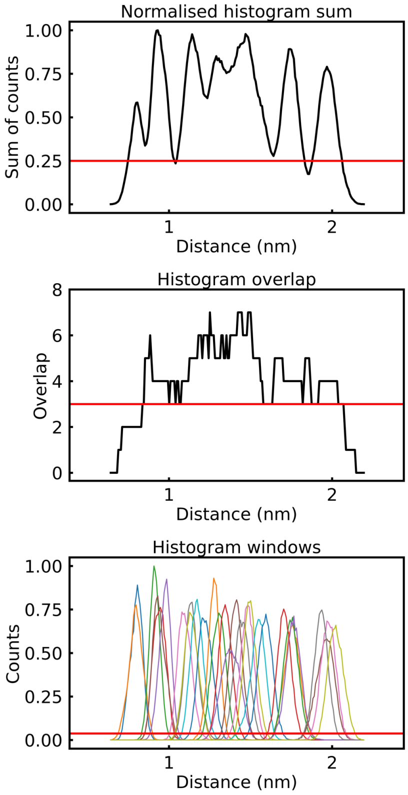


#### RHBDL2

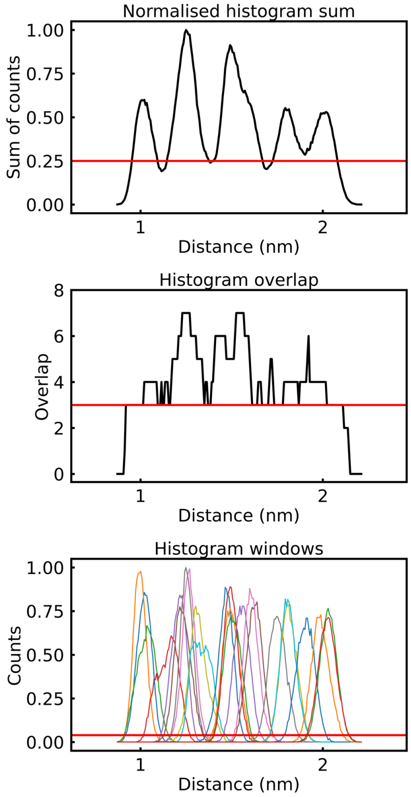


#### RHBDL3

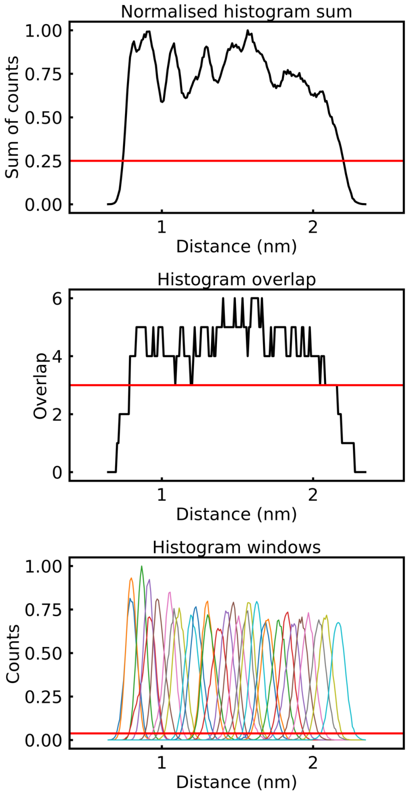


#### RHBDL4

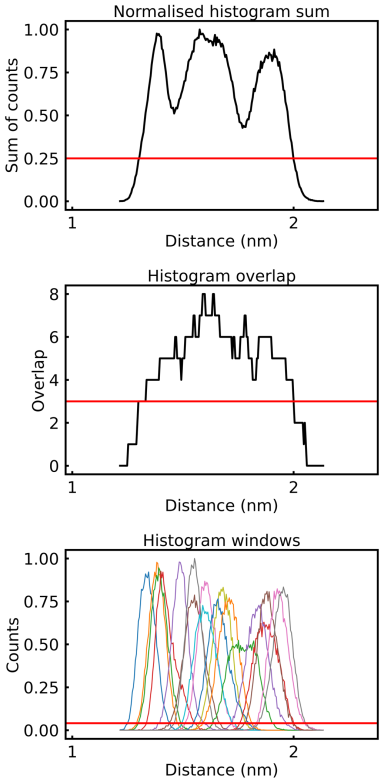

### Figure S14

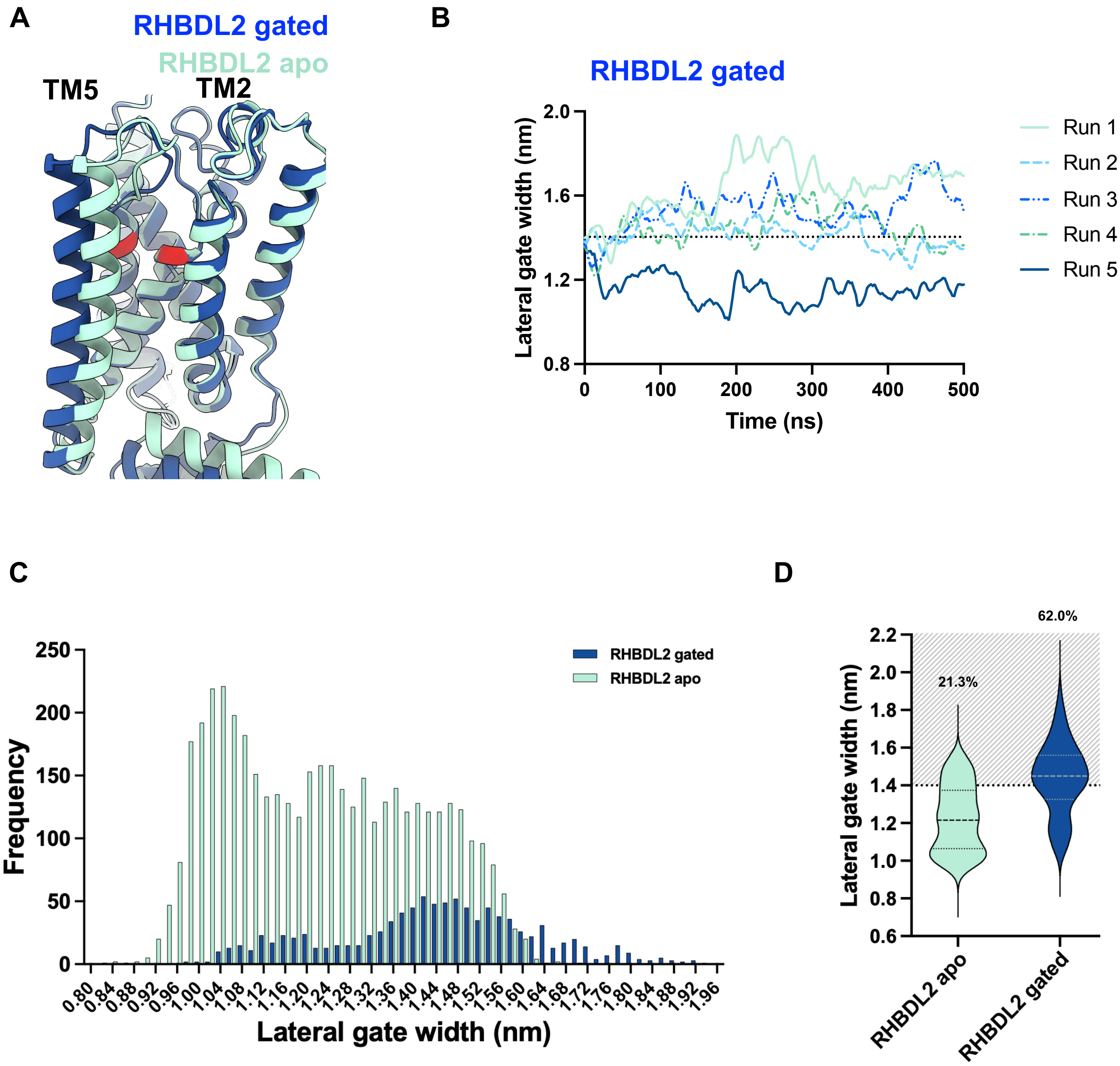

A
RHBDL2 gated
RHBDL2 apo
TM5
TM2
B
RHBDL2 gated
Lateral gate width (nm)
Time (ns)
Run 1
Run 2
Run 3
Run 4
Run 5
C
Frequency
Lateral gate width (nm)
RHBDL2 gated
RHBDL2 apo
D
Lateral gate width (nm)
21.3%
62.0%
RHBDL2 apo
RHBDL2 gated

### Figure S15

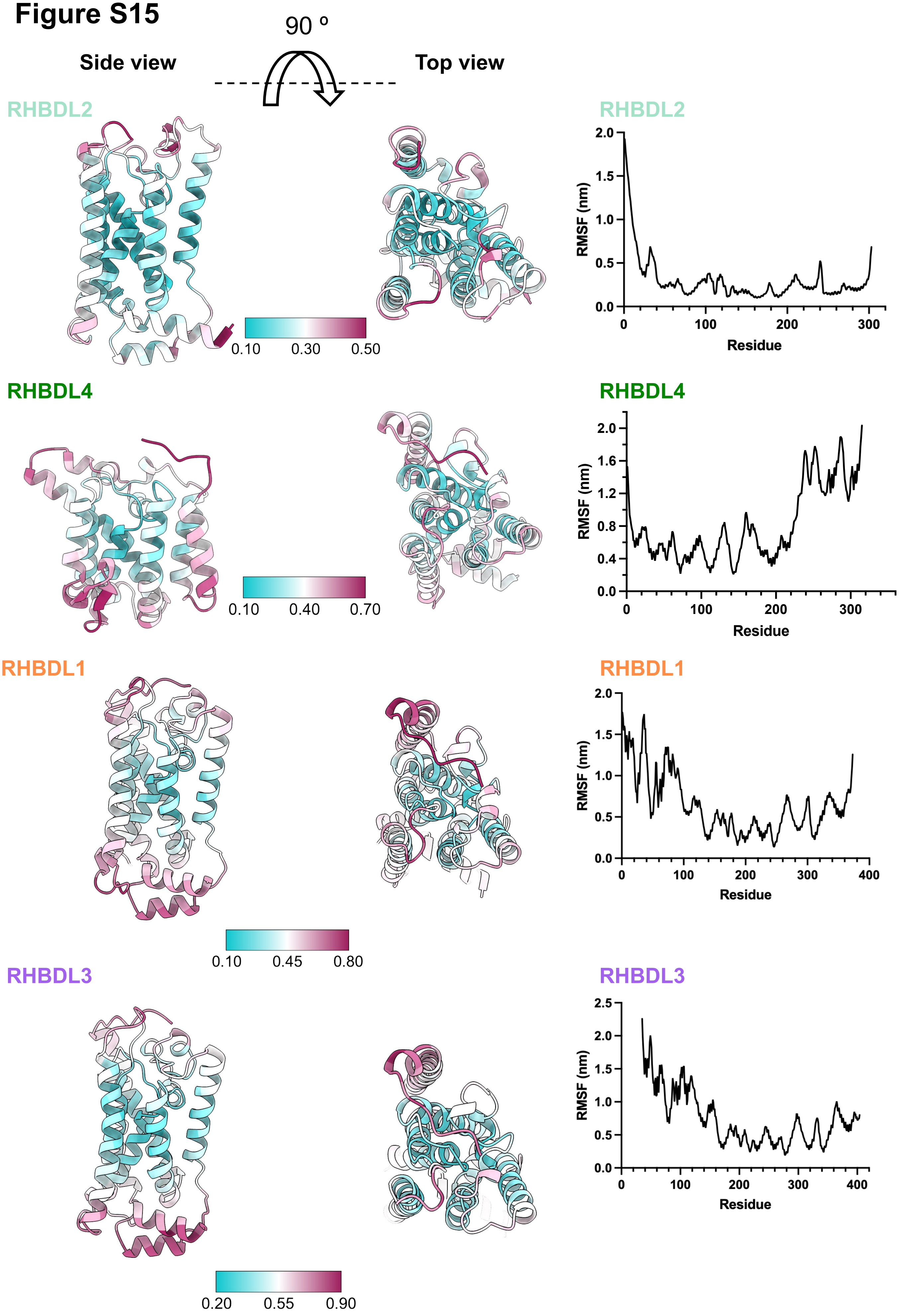

90 º
Side view
Top view
RHBDL2
0.10
0.30
0.50
RHBDL2
RMSF (nm)
Residue
RHBDL4
0.10
0.40
0.70
RHBDL4
RMSF (nm)
Residue
RHBDL1
0.10
0.45
0.80
RHBDL1
RMSF (nm)
Residue
RHBDL3
0.20
0.55
0.90
RHBDL3
RMSF (nm)
Residue

### Figure S16

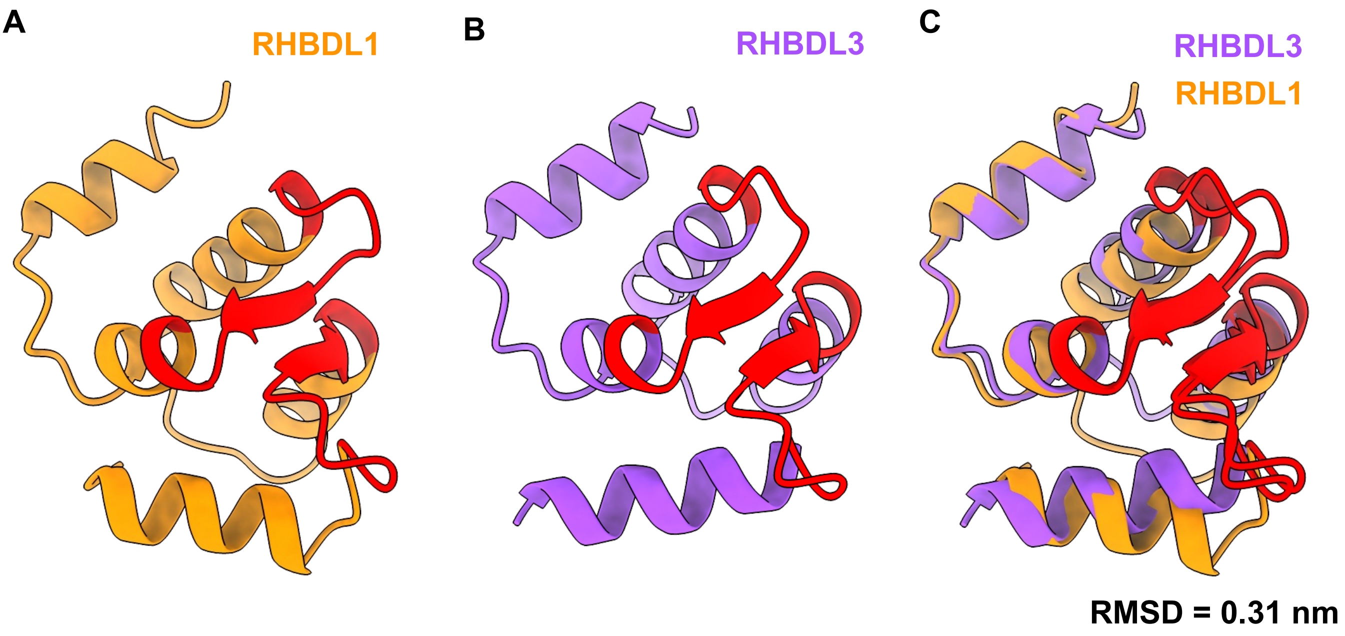


D **EF hands consensus motif: DxDxxGxIxxxE**

```
RHBDL1      MDRSSLLQLIQEQQLDPENTGFIGADTFTGLVHSHELPLDPAKLDML  47
RHBDL3      ---PEDHWKVLFDQFDPGNTGYISTGKFRSLLESHSSKLDPHKREVL  78
                .    :  :*:** ***:*.:..* .*:.**.  *** * ::*

RHBDL1      VALAQSNEQGQVCYQELVDLISSKRSSSFKRAIANGQ  84
RHBDL3      LALADSHADGQIGYQDFVSLMSNKRSNSFRQAILQGN  115
            :***:*: :**: **::*.*:*.***.**::** :*:
```

### Supplementary Table 1

| Model | AF2 | Chai-1 | Boltz-2 |
|---|---|---|---|
| **RHBDL1** | 0.744 | 0.766 | 0.774 |
| **RHBDL2** | 0.840 | 0.837 | 0.809 |
| **RHBDL3** | 0.710 | 0.767 | 0.724 |
| **RHBDL4** | 0.636 | 0.698 | 0.674 |
| **PARL** | 0.853 | 0.857 | 0.815 |
| **EcGlpG** | 0.659 | 0.750 | 0.672 |

### Supplementary Table 2

| Model | MolProbity score | Ramachandran outliers (%) | Clash score | Poor rotamers (%) | Average local score | C-beta z-score | Interaction z-score | Packing z-score | QMEAN4 Z-score | QMEAN6 Z-score | Torsion z-score |
|---|---|---|---|---|---|---|---|---|---|---|---|
| **RHBDL1** | 1.49 | 1.08 | 2.2 | 1.60 | 0.814 | -2.339 | 1.476 | 1.338 | -2.368 | -2.247 | -2.584 |
| **RHBDL2** | 1.10 | 1.33 | 1.04 | 0.40 | 0.831 | -0.424 | 1.378 | 1.838 | -2.504 | -2.762 | -3.248 |
| **RHBDL3** | 1.33 | 0.82 | 2.03 | 0.63 | 0.774 | -1.603 | 1.280 | -1.164 | -4.047 | -3.520 | -3.674 |
| **RHBDL4** | 1.83 | 4.79 | 1.8 | 2.60 | 0.736 | -0.443 | 0.288 | 0.449 | -2.637 | -2.472 | -2.837 |
| **PARL** | 0.66 | 0.00 | 0.46 | 0.43 | 0.794 | -1.354 | 1.175 | 1.011 | -2.766 | -2.663 | -3.050 |
| **EcGlpG** | 1.21 | 2.55 | 1.58 | 0.90 | 0.831 | -2.494 | 0.331 | 1.343 | -1.395 | -1.625 | -1.500 |
| **2IC8** | 2.59 | 0.56 | 15.1 | 6.90 | N/A | N/A | N/A | N/A | N/A | N/A | N/A |
| **2NRF** | 3.51 | 3.72 | 40.14 | 10.60 | N/A | N/A | N/A | N/A | N/A | N/A | N/A |
